# Discovery and chemical biology of CDK8 inhibitors reveals insights for kinase inhibitor development

**DOI:** 10.1101/2025.11.06.686927

**Authors:** Franck Borel, Yann Vai Le Bihan, Béatrice Foll-Josselin, Thomas Robert, Julien Duez, Lucie Angevin, Elisabeth Simboeck, Susana Prieto, Elsie Hodimont, Jean-Luc Ferrer, Liliana Krasinska, Sandrine Ruchaud, Daniel Fisher

## Abstract

Protein kinase inhibitors are a key class of targeted cancer therapies, but finding compounds that show on-target efficacy with minimal toxicity remains challenging. CDK8 and CDK19 are paralogous kinases that regulate Mediator complex to promote transcriptional responses to extracellular signals and have also been reported to facilitate genome replication. Preclinical studies show that CDK8/19 inhibitors may have therapeutic benefit in several cancers, but whether effective inhibition of Mediator kinases is cytotoxic is debated. Here, we present a multi-modal approach for discovery of specific kinase inhibitors, and find that most CDK8/19 inhibitors show off-target toxicity and/or insufficient on-target engagement in cells. We describe several novel high affinity CDK8/19 inhibitors, report that loss of CDK8/19 activity is not cytotoxic but reduces cell proliferation without affecting DNA replication, and provide evidence that most inhibitors can bind in type I or type II modes. Our approach is applicable to any kinase inhibitor development program.

## Introduction

Protein kinase inhibitors have become standard-of-care single-agent therapies for many cancers, including previously intractable types such as non-small cell lung cancer (*e.g.* ^1^). Kinase inhibitors may also show significant benefits when combined with other treatments (*e.g.* ^2^). Most current widely-used inhibitors target growth factor receptor tyrosine kinases such as EGFR, ALK, and c-ABL. Resistance, driven by mutations or activation of alternative signalling pathways, along with toxicity, remain significant challenges that direct further kinase inhibitor development ^3^. Targeting downstream serine/threonine kinases that more directly control cell proliferation has also been extensively explored. Cyclin-dependent kinases (CDKs) are a family of 20 such kinases. The CDK1/CDK2 subfamily directly triggers onset of S-phase and mitosis ^4^, while CDK4 and CDK6 indirectly promote cell cycle progression by inactivating the repressor of cell cycle-regulated transcription, RB. Thus, CDK inhibitors have long been seen as potential cancer therapeutic agents. Although early trials with pan-CDK inhibitors such as flavopiridol were limited by excessive toxicity, more selective molecules show considerable promise. As a result of unprecedented benefits in clinical trials combined with limited toxicity, specific inhibitors of CDK4/6 have been approved in combination with anti-oestrogen drugs for advanced metastatic HER2+ breast cancers, and they are currently undergoing trials for many other cancer types ^5,6^.

Another class of CDKs directly controls transcription. This includes the highly conserved subfamily composed of CDK8 and CDK19, which share an essentially identical kinase domain. Both CDK8 and CDK19 bind cyclin C, and each of these complexes interacts with MED12 and MED13 (or their paralogues, MED12L and MED13L), to form the kinase module of the Mediator transcriptional regulator. Whereas Mediator is essential and promotes the expression of all genes transcribed by RNA polymerase II ^7^, the kinase module is non-essential for cell survival and proliferation in most cell types and in most species ^8–15^. Consequently, the physiological functions of CDK8 and CDK19 remain unclear. However, CDK8 and CDK19 have been widely implicated in oncogenic and immune-regulatory transcriptional programmes ^10,16–26^. CDK8 itself has been attributed oncogenic activity in a subclass of colorectal cancers in which Wnt pathway signalling is deregulated ^27^, although this has not yet been unequivocally confirmed *in vivo* and genetic models of intestinal cancer have not revealed a strong involvement of CDK8 to date ^12,13,28^. CDK8 or CDK19 have also been identified as useful therapeutic targets in Acute Myeloid Leukaemia (AML) where they appear to dampen rather than promote gene expression, in particular at super-enhancers ^21^. CDK8 and/or CDK19 may also have kinase-independent roles in cancer: CDK8 deletion, but not inhibition, reduced survival of B-cell Acute Lymphoblastic Leukaemia (B-ALL) cells in pre-clinical mouse models ^29^. *In vitro* experiments with cultured cell lines suggest that CDK8 and CDK19 are interchangeable and act as amplifiers of signal-regulated transcription ^14^. As such, their specific inhibition may not be excessively toxic, but might limit the deregulated transcription of cancer cells to restore a less aggressive phenotype. Consistent with a role of transcriptional amplifiers, CDK8 and CDK19 inhibition has proven beneficial in reversing resistance to chemotherapies or targeted therapies in preclinical models of breast and prostate cancer ^30–32^, and in increasing effectiveness of anti-tumour immune responses ^24,26,33^.

Mechanistically, CDK8 and CDK19 can phosphorylate components of the Mediator complex, disrupting its binding to the CDK module, thus allowing Mediator to interact with the transcription pre-initiation complex ^34^. They can also directly phosphorylate SMAD, NOTCH and STAT1 transcription factors to regulate their activity ^10,18,33,35^, as well as the C-terminal domain of RNA polymerase to stimulate transcriptional elongation ^16,36^. Yet the mechanisms by which CDK8 and CDK19 phosphorylate specific targets and control a limited subset of genes have not yet been elucidated, and how their activity limits transcription stimulated by super-enhancers is unknown.

Finally, CDK8 and/or CDK19 appear to have cellular effects that are independent of transcription, suggesting that they may affect processes deregulated in cancer by multiple mechanisms. For example, they have been reported to promote completion of DNA replication prior to mitosis by direct binding to the replication factor MTBP ^37^, and their inhibition affects both splicing and metabolism in a cell type-specific manner ^38^.

Despite the uncertainty remaining about their precise mechanisms of action and physiological functions, at least 35 lead small molecule inhibitors of CDK8 and CDK19 have been described to date and characterised to a greater or lesser degree (Figure 1 and Table S1). Some of these inhibitors have shown benefit in pre-clinical models of different cancers (Table S1). Over 30 structures of CDK8-cyclin C with inhibitors are available (Table S2). All prevent ATP-binding. As with other kinase inhibitors, most have been assigned to one of two different classes, type I (ATP-competitive) and type II inhibitors. The distinction concerns the inhibitor-bound position of the activation loop orienting the essential aspartate-173 within the DMG motif, conserved in all eukaryotic protein kinases, which in the active conformation forms contacts with all three phosphate groups of ATP ^39^. Crystal structures of inhibitors bound to CDK8/CDK19 where this triplet is present in the active conformation (“DMG-in”) designate them as type I inhibitors, whereas type II inhibitors are those in which the crystal structures reveal the DMG oriented away from the active site (“DMG-out”), exposing a deep hydrophobic pocket that forms additional contacts with the inhibitor ^40–42^. The extent to which inhibitors employ these type I/II binding modes in living cells, and their relevance for physiological effects of the inhibitors, remain unknown.

**Fig. 1.**
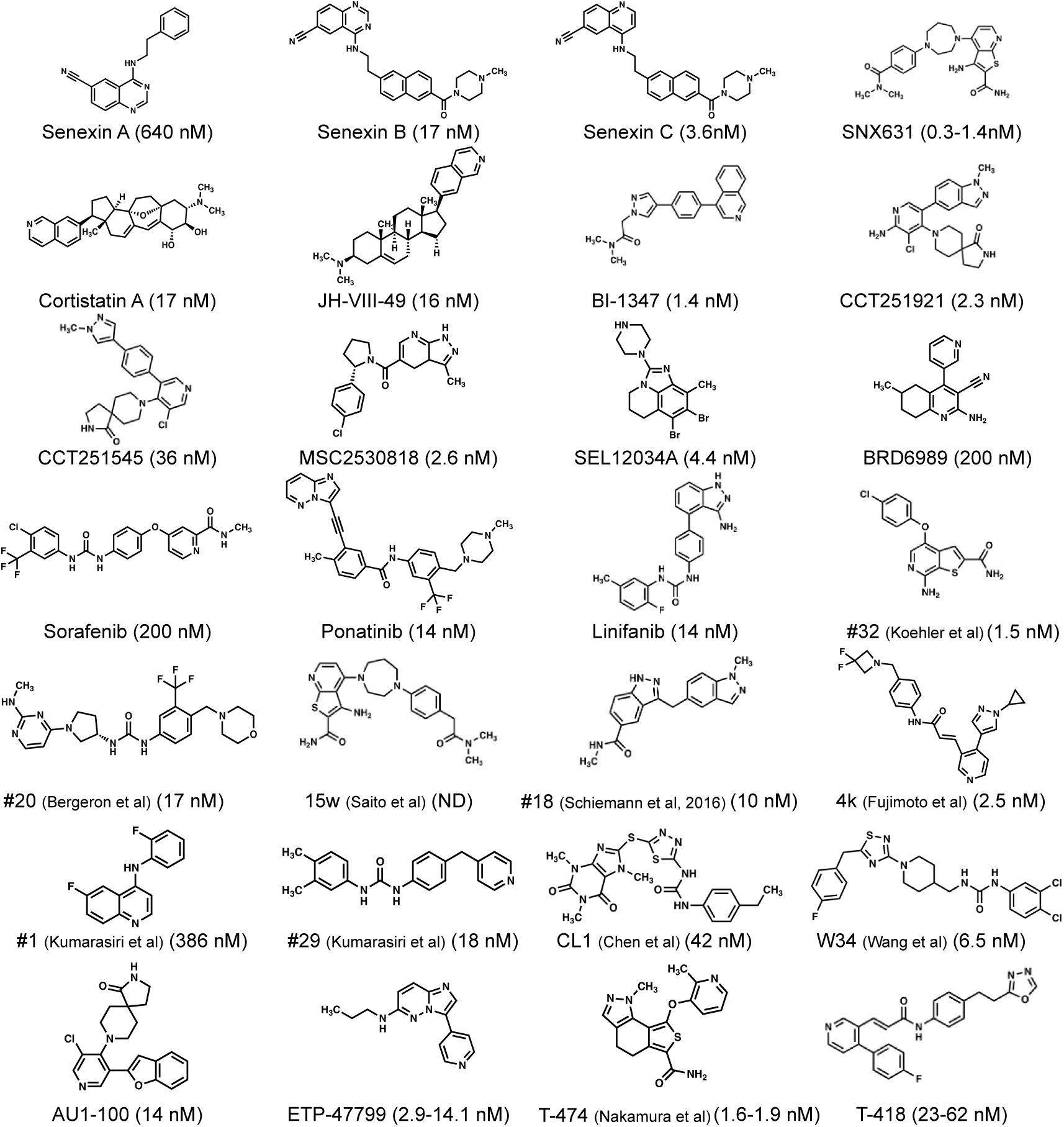
Structures of CDK8/19 inhibitors published to date. A single molecule with best characteristics is chosen from each publication (see Table S1 for futher detail). IC_50_ values are indicated.

To date, specific on-target engagement with on-target toxicity has not been shown for most CDK8/CDK19 inhibitors published. For example, most CDK8/CDK19 inhibitors have been found to reduce proliferation of specific cell lines (Table S1), yet some type I inhibitors have also revealed equivalent toxicity in cells lacking CDK8 and CDK19 ^43,44^, demonstrating that it is not CDK8/19-dependent. This is consistent with genetic studies that have disrupted both CDK8 and CDK19 and revealed that most mammalian cells can proliferate normally without either paralogue ^12–15^. *In vivo* toxicity in some animal models was argued, though not proven, to be on-target (*i.e.* CDK8/CDK19-dependent) ^45^, yet other CDK8/CDK19 inhibitors do not show equivalent animal toxicity and have markedly different effects on the transcriptome, implying that toxicity and transcriptome effects are related and may be off-target ^46^. Some inhibitors have limited activity in cell-based assays ^41,47^, and most have either not been confirmed to bind to CDK8 or CDK19 in cells ^41^, or target engagement in cells has not been tested. Overall, the absence of a clear specific cellular readout of CDK8/CDK19 activity has proven problematic, as transcriptional effects of inhibitors vary widely and are cell line-dependent, while no *in vivo* phosphorylation sites, including the routinely used phospho-STAT1-S727, are completely specific to CDK8/CDK19 ^46^. Knowledge of specificity is critical to account for toxicity, yet the kinase specificity of inhibitors has been assessed in many different ways, including ligand displacement assays such as Lanthascreen^TM^ and Kinomescan^TM^, that may give different results from *in vitro* phosphorylation assays involving CDK8-cyclin C and a substrate.

Such difficulties are far from being unique to CDK8/CDK19 inhibitors. A variety of common shortcomings have plagued cancer drug target development, including on-target versus off-target effects and the mis-interpretation of correlation as causation, often as a result of over-reliance on single assays and the inadequate use of appropriate controls ^48^.

Here, by addressing the difficulties highlighted above, we present a rational unbiased approach for discovery and analysis of CDK8/CDK19 inhibitors. We report the identification of several new potent and specific CDK8/CDK19 inhibitors. We find that cellular toxicity of most CDK8/CDK19 inhibitors is independent of target inhibition, in line with recent studies showing that most common cancer drugs act via off-target toxicity ^49^. We also find evidence that CDK8 and CDK19 are required for optimal proliferation of primary cells but are dispensable for highly efficient DNA replication. Finally, molecular modelling shows that most, if not all CDK8i, can exhibit both type I and type II binding modes; thus, their classification as type I or type II might not be accurate. In summary, we propose an effective approach to address several key general limitations for kinase inhibitor development.

## Results

The first CDK8/CDK19 inhibitors were identified serendipitously, for example through characterisation of natural molecules with biological activity ^50^, or from a variety of cell-based phenotypic screens ^23,51,52^, and thus were not designed to target CDK8/19 specifically. However, target-based approaches including medium-throughput screens of several thousand molecules for CDK8 ligands ^24^ and virtual screening of several hundred thousand molecules against the CDK8 structure ^53,54^ have also been used successfully. We reasoned that the latter strategy has unexploited potential as an efficient first-line approach for inhibitor discovery. Known CDK8/CDK19 inhibitors score well in molecular docking analyses and libraries of over 10^7^ existing molecules are available. We generated a structure for virtual screening based on the crystal structure (PDB: 3RGF) of CDK8-cyclin C bound to Sorafenib ^40^, and modelling in the activation loop (absent from this structure) based on the active CDK2-cyclin A-ATP structure ^55^. Docking of Sorafenib (Figure S1) recapitulated the binding mode seen in the crystal structure ^40^ and gave a docking score (DS) of -13.2, while the DS for Senexin A was -10.1, confirming the accuracy of the model. We then performed virtual screening with the curated Zinc database ^56^, that now includes essentially all commercially available molecules. We used Glide ^57^ in High Throughput Virtual Screening (HTVS) mode ^58^ to screen 21.5 million molecules of the Zinc12 database ^56^. Compounds were ranked by DS and the best were re-screened iteratively in Standard Precision (SP) and Extra Precision (XP) ^59^ mode.

This identified 229 molecules with a very good (XP DS <-12) docking score for CDK8 binding, corresponding to a low free energy in the bound state that typically correlates with nanomolar affinity. Finally, we re-performed the docking using induced fit parameters, which confirmed the excellent docking scores (Dataset S1). We then acquired the 99 best candidate molecules for biochemical testing in a secondary screen. We tested each inhibitor at a concentration of 10 µM in radiometric protein kinase assays using 90 nM purified recombinant CDK8-cyclin C, the RBER-IRStide peptide substrate and ^33^P-ATP. 36 compounds showed more than 50% inhibition of CDK8-cyclin C in these conditions (Figure S2). We re-screened all of these at the same concentration against a panel of in-house purified protein kinases that are frequently found as off-targets of current CDK8/CDK19 inhibitors (Table S1). This set comprised CDK5-p25, CDK2-cyclin A, DYRK1A, GSK3-α/β, CLK1, PIM1 and Haspin. Only three compounds inhibited any of these kinases at >50%, all of which inhibited CLK1 (Table 1). In comparison, the well-characterised CDK8 inhibitor Senexin A partially inhibited 7 of the 8 potential off-targets and 3 of them at over 50%. We next performed further assays with our compounds at a range of concentrations to determine the IC_50_, which varied from 5 nM (compound #82) to 7 µM (compound #21), with 26 inhibitors (72%) having sub-micromolar IC_50_ and 6 (16%) of them showing IC_50_ below 100 nM (Figure 2). This highlights the efficiency of a virtual screening approach in identifying specific inhibitors.

**Fig 2.**
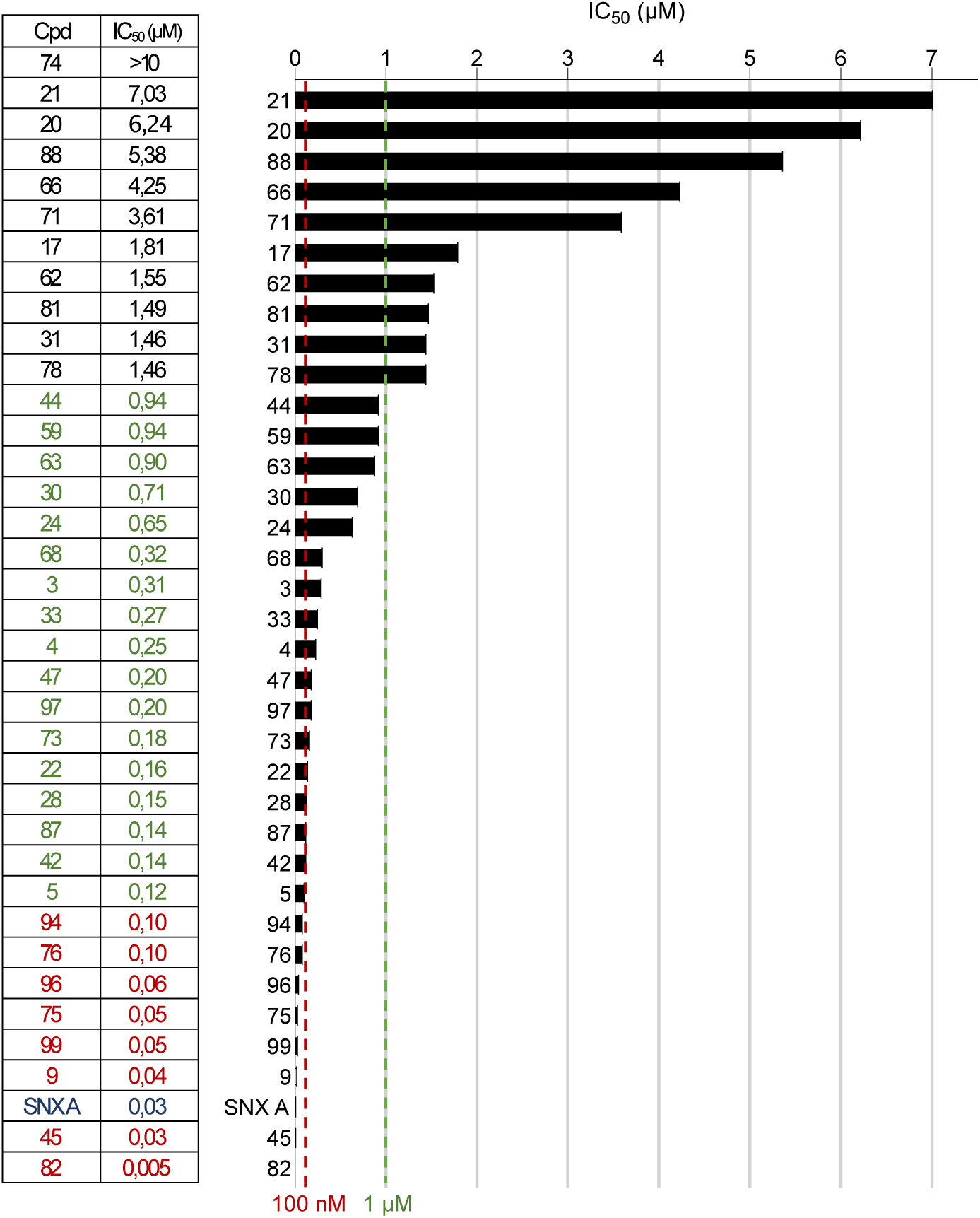
IC_50_ evaluation of selected compounds on CDK8 activity. Thirty six selected compounds and Senexin A (SNX A) were tested in dose-response on CDK8/Cyc C in kinase assays. IC_50_ is expressed in µM and calculated from dose-response curves (each curve was performed in duplicate). The graph represents the distribution of IC_50_ obtained, with two threshold lines: at 100 nM (red line) and 1 µM (green line).

**Table 1.** Selectivity of screening hits in kinase and cell viability assays. Left, percentage of kinase residual activities in the presence of 10 μM compound concentration. Kinase inhibitory activities were assayed in duplicate in the presence of 15 μM ATP. Right, effect of selected compounds on cancer and non-cancer cell viability. Cells were incubated for 48h with increasing doses of each compound (up to 50 μM). Cell viability was determined by MTS assay in triplicate, and EC_50_ (μM) were calculated from the dose-response curves.

| *cmpd* | *Selectivity, % residual activity (10 μM)* |  |  |  |  |  |  |  | *Viability test (EC<sub>50</sub>, μM)* |  |  |  |
| --- | --- | --- | --- | --- | --- | --- | --- | --- | --- | --- | --- | --- |
| | CDK8/<br>cyclin C | CDK5/<br>P25 | CDK2/<br>cyclin A | CLK1 | DYRK1A | GSK3<br>-$\alpha/\beta$ | HASPIN | PIM1 | RPE | HCT116 | SHSY-5Y | U2OS |
| 3 | 19 | 121 | 83 | 72 | 65 | 110 | 90 | 100 | 34 | 45 | >50 | 36 |
| 4 | 18 | 93 | 69 | 107 | 70 | 115 | 100 | 118 | 40 | 35 | >50 | >50 |
| 5 | 14 | 111 | 67 | 100 | 78 | 115 | 103 | 44 | 12 | 20 | >50 | 12 |
| 9 | 10 | 107 | 78 | 111 | 80 | 119 | 94 | 71 | >50 | >50 | >50 | >50 |
| 17 | 18 | 108 | 85 | 97 | 75 | 118 | 122 | 81 | 21 | 49 | >50 | 37 |
| 20 | 43 | 106 | 71 | 65 | 86 | 114 | 95 | 82 | >50 | >50 | >50 | >50 |
| 21 | 46 | 169 | 146 | 154 | 97 | 123 | 161 | 130 | 12 | >50 | >50 | >50 |
| 22 | 10 | 115 | 76 | 151 | 83 | 113 | 96 | 85 | 10 | 29 | 41 | 14 |
| 24 | 32 | 115 | 76 | 71 | 54 | 112 | 98 | 114 | 17 | 14 | >50 | 23 |
| 28 | 9 | 119 | 75 | 109 | 67 | 106 | 89 | 87 | 12 | >50 | >50 | 31 |
| 30 | 11 | 119 | 57 | 112 | 78 | 119 | 97 | 67 | 14 | >50 | >50 | 24 |
| 31 | 16 | 98 | 79 | 94 | 73 | 121 | 107 | 62 | 12 | >50 | >50 | 35 |
| 33 | 45,4 | 92 | 70 | 93 | 78 | 99 | 107 | 67 | >50 | >50 | >50 | >50 |
| 42 | 6 | 93 | 66 | 10 | 89 | 91 | 96 | 58 | 46 | >50 | >50 | 28 |
| 44 | 14 | 103 | 76 | 141 | 100 | 119 | 118 | 68 | 25 | 38 | 34 | 21 |
| 45 | 6 | 115 | 82 | 143 | 73 | 88 | 144 | 127 | 43 | 19 | 8 | 9 |
| 47 | 5 | 107 | 81 | 109 | 75 | 115 | 118 | 89 | >50 | >50 | >50 | >50 |
| 59 | 22 | 110 | 94 | 131 | 104 | 105 | 116 | 119 | >50 | >50 | >50 | >50 |
| 62 | 20 | 98 | 103 | 106 | 101 | 92 | 107 | 76 | >50 | >50 | >50 | >50 |
| 63 | 11 | 108 | 100 | 114 | 90 | 104 | 105 | 95 | >50 | >50 | >50 | >50 |
| 66 | 46 | 117 | 77 | 112 | 70 | 92 | 103 | 82 | >50 | >50 | >50 | >50 |
| 68 | 25 | 103 | 92 | 116 | 83 | 95 | 93 | 91 | >50 | >50 | >50 | >50 |
| 71 | 30 | 111 | 103 | 121 | 110 | 110 | 91 | 93 | 21 | >50 | >50 | >50 |
| 73 | 8 | 85 | 82 | 103 | 76 | 112 | 100 | 103 | >50 | >50 | >50 | >50 |
| 74 | 39 | 92 | 97 | 109 | 99 | 87 | 101 | 89 | >50 | >50 | >50 | >50 |
| 75 | 8 | 101 | 106 | 99 | 89 | 104 | 123 | 95 | >50 | >50 | >50 | >50 |
| 76 | 11 | 89 | 73 | 37 | 80 | 97 | 116 | 109 | >50 | >50 | >50 | >50 |
| 78 | 45 | 93 | 75 | 96 | 74 | 92 | 111 | 78 | 41 | >50 | 44 | 31 |
| 81 | 21 | 109 | 86 | 110 | 73 | 90 | 109 | 88 | >50 | >50 | >50 | >50 |
| 82 | 19 | 109 | 120 | 81 | 134 | 105 | 111 | 89 | >50 | 31 | 24 | 10 |
| 87 | 21 | 108 | 94 | 66 | 84 | 111 | 92 | 79 | >50 | >50 | >50 | 45 |
| 88 | 48 | 98 | 109 | 80 | 74 | 100 | 108 | 77 | >50 | >50 | >50 | >50 |
| 94 | 20 | 104 | 101 | 71 | 105 | 94 | 98 | 92 | >50 | >50 | >50 | >50 |
| 96 | 19 | 104 | 100 | 58 | 91 | 91 | 102 | 83 | >50 | >50 | >50 | 41 |
| 97 | 33 | 116 | 108 | 22 | 48 | 99 | 103 | 77 | 7 | >50 | >50 | >50 |
| 99 | 15,8 | 65 | 84 | 25 | 67 | 79 | 91 | 95 | 7 | 29 | 12 | 17 |
| SNX A | 15 | 57 | 62 | 34 | -7 | 72 | ND | 78 | 41 | >50 | >50 | 48 |

CDK8/CDK19 inhibitors have been claimed to have both on-target and off-target inhibitory activity for cell viability in different cell lines. As ascertained by compilation of genome-wide CRISPR-Cas9 knockout screens in over 1100 cancer cell lines by the DepMap consortium ^60^ (https://depmap.org), ablation of CDK8 or CDK19 alone has negligible effects on growth in the vast majority of cell lines (Figure S3A). We similarly analysed all CDKs and several kinases often found as off-targets of CDK8/19 inhibitors, as well as the other components of the Mediator kinase module (Figure S3B). All cell lines depend on the mitotic kinase CDK1, while ablation of other cell cycle CDKs, CDK2, CDK4 and CDK6 strongly affects most cell lines. Of the other known “transcriptional” CDKs, CDK7 and CDK9 are essential in all cell lines and CDK12 in the vast majority, in contrast to CDK8 and CDK19. Only just over half of cell lines are affected by the ablation of MED12, while ablation of the single activating subunit, cyclin C (which activates both CDK8 and CDK19) reduces cell growth in about 25% of cell lines, and MED13 dependency was less than 5% of cell lines. The most common off-targets of known CDK8/19 inhibitors (Table S1) are largely non-essential. This analysis suggests that on-target toxicity of inhibitors is unlikely to be systematic, and will probably be cell line-dependent. It also suggests that the highest risk for toxic off-target effects are other CDKs.

To assess this experimentally, we determined EC_50_ (concentration that inhibits growth, or viability, by 50% compared to vehicle) of our 36 inhibitors and Senexin A against three human cancer cell lines with different driver mutations (Table 1). We used the MYCN-amplified, p53-mutant neuroblastoma line SH-SY5Y; the RB1-mutant, p16-mutant and MDM2-amplified U2OS osteosarcoma line; and the beta catenin-mutant, SMAD4-mutant, KRAS^G13D^-mutant, mismatch repair-deficient but p53-proficient HCT-116 colorectal carcinoma line. We also used non-transformed hTERT immortalised retinal pigment epithelial (RPE-1) cells. As assessed by a Spearmann rank correlation test, EC_50_ of the 37 inhibitors did not significantly correlate with IC_50_ for CDK8-cyclin C for any cell line. RPE-1 cells were sensitive to more compounds than the cancer cell lines. Five compounds were toxic (EC_50_ < 50 µM) against SH-SY5Y cells, of which only three (#22, #45, #82) have low EC_50_ for CDK8-cyclin C; these three compounds all had multiple cell line toxicities. In contrast, two compounds (#9 and #75) had low IC_50_ for CDK8-cyclin C but no detectable toxicity against any cell line. There are several possible reasons for the lack of correlation between IC_50_ for CDK8-cyclin C and cellular toxicity: it could be that toxicity is not due to inhibition of CDK8-cyclin C but non-specific effects, or that it is on-target yet inhibitors that have low IC_50_ for CDK8-cyclin C but no obvious toxicity do not efficiently reach their target in cells, either through permeability problems or sequestration by other cellular components.

To explore this further, we focused on a subset of seven inhibitors (#9, #22, #45, #47, #75, #76, #82; Figure 3) with similar IC_50_ for CDK8-cyclin C (5-200 nM) but different toxicity profiles. Comparing induced fit docking scores for each inhibitor against CDK8-cyclin C and CDK2-cyclin A revealed large differences, explaining their specificity for CDK8 (Figure 3). We next assessed toxicity of these seven inhibitors and Senexin A against primary mouse embryonic fibroblasts (MEFs), permitting comparison with MEFs genetically inactivated for CDK8 and CDK19. As with the four human cell lines tested, compounds #22, #45, #82 and Senexin A at 10 µM significantly inhibited 5-day growth and colony formation of wild-type MEFs (Figure 4A, B), with Senexin A proving the most toxic, while compounds #9, #47, #75 and #76 had little effect, confirming that cellular effects are reproducible across different human and murine cell lines. To assess the extent to which *in vitro* toxicity is dependent on target inhibition, we performed similar experiments on immortalised MEFs lacking Mediator kinases (Angevin *et al.,* unpublished results) derived from *Rpb1::Cre^ERT2^;Cdk8^lox/lox^* mice ^13^. In these cells, tamoxifen treatment for 14 days induced recombination and loss of exon 2 of the *Cdk8* gene, removing the essential catalytic lysine-52 and causing a frameshift mutation resulting in truncation of 90% of the protein ^13^. We also tested inhibitors on MEFs in which CDK19 alone was ablated by CRISPR-Cas9, and MEFs lacking both CDK8 and CDK19. The double mutant did not show a loss of viability (Figure 4C). In all cases, the reduction of growth in the presence of the inhibitors #22, #45, #82 and Senexin A was not alleviated by either of the single mutants, and only partly by the double mutation (Figure 4C). This suggests that the majority of cellular toxicity shown by inhibitors is independent of target inhibition, but it does not rule out the possibility that specific target inhibition is toxic. We next tested whether the compounds showing significant toxicity, #22, #45 and #82, inhibit phosphorylation of STAT1-S727 in cells, which is thought to be largely dependent on CDK8. We also compared with Senexin A and the more recent compound Senexin B. In MEFs, #82 at 10µM attenuated the interferon-γ-mediated induction of STAT1-S727ph, but was not effective at 1µM, unlike both Senexin A and Senexin B. #22 and #45 had no effect at any concentration (Figure 4D). We obtained similar results when treating colonospheres of HCT-116 cells (Figure 4E). This provides further evidence that the toxicity of #22 and #45 is not dependent on the efficient inhibition of CDK8/CDK19, while #82 appears to be an effective inhibitor yet has some off-target toxicity.

**Fig. 3.**
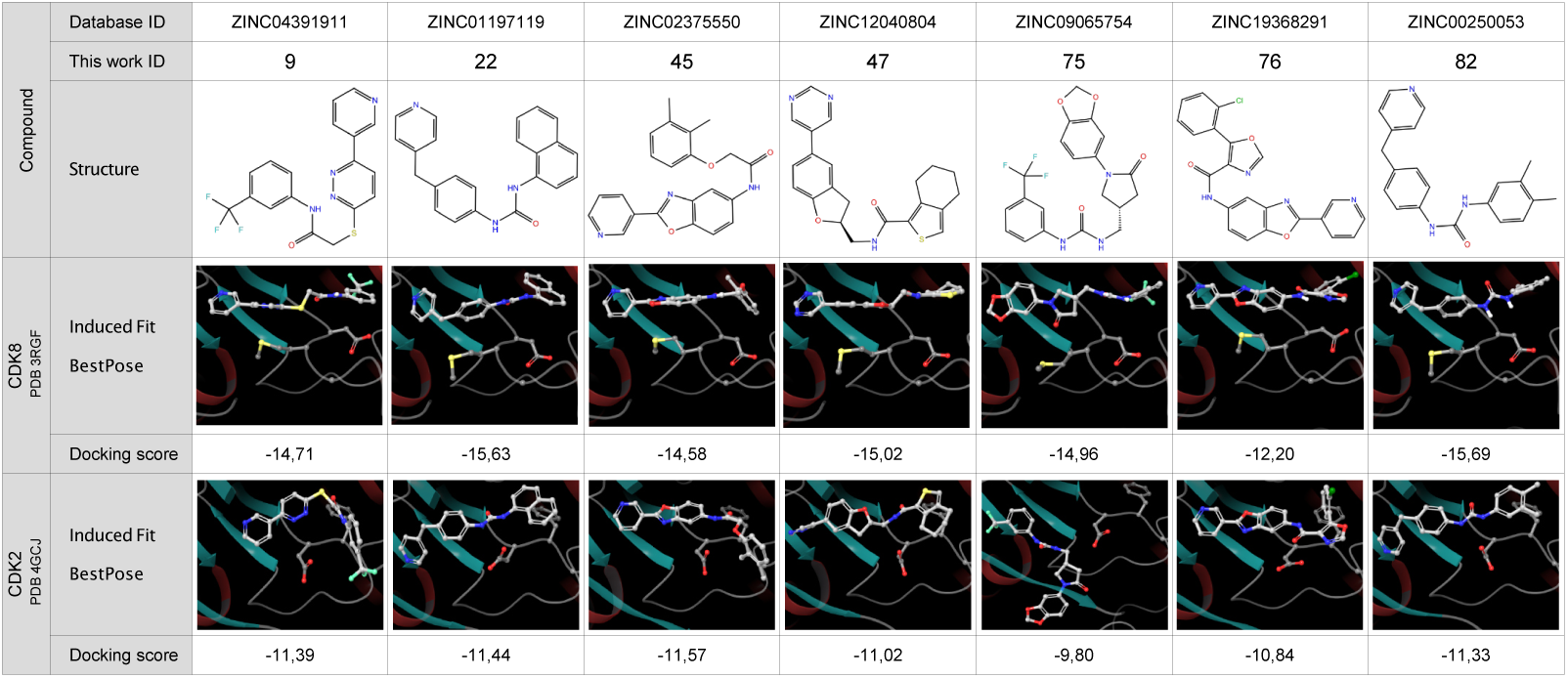
Structures and in silico docking of selected compounds. Induced fit docking experiments were performed using either CDK8 (PDB: 3RGF) or CDK2 (PDB: 4GCJ) structures as protein target. Upper panels: ID and structural formula of the seven selected compounds. Lower panels: 3D arrangement of the best docking pose (white carbon atoms) into CDK8 or CDK2 active site with its corresponding docking score. Amino acids from the CDK8 DMG loop are represented in ball and stick; corresponding residues (DFG) in CDK2 structure are also highlighted.

**Fig. 4.**
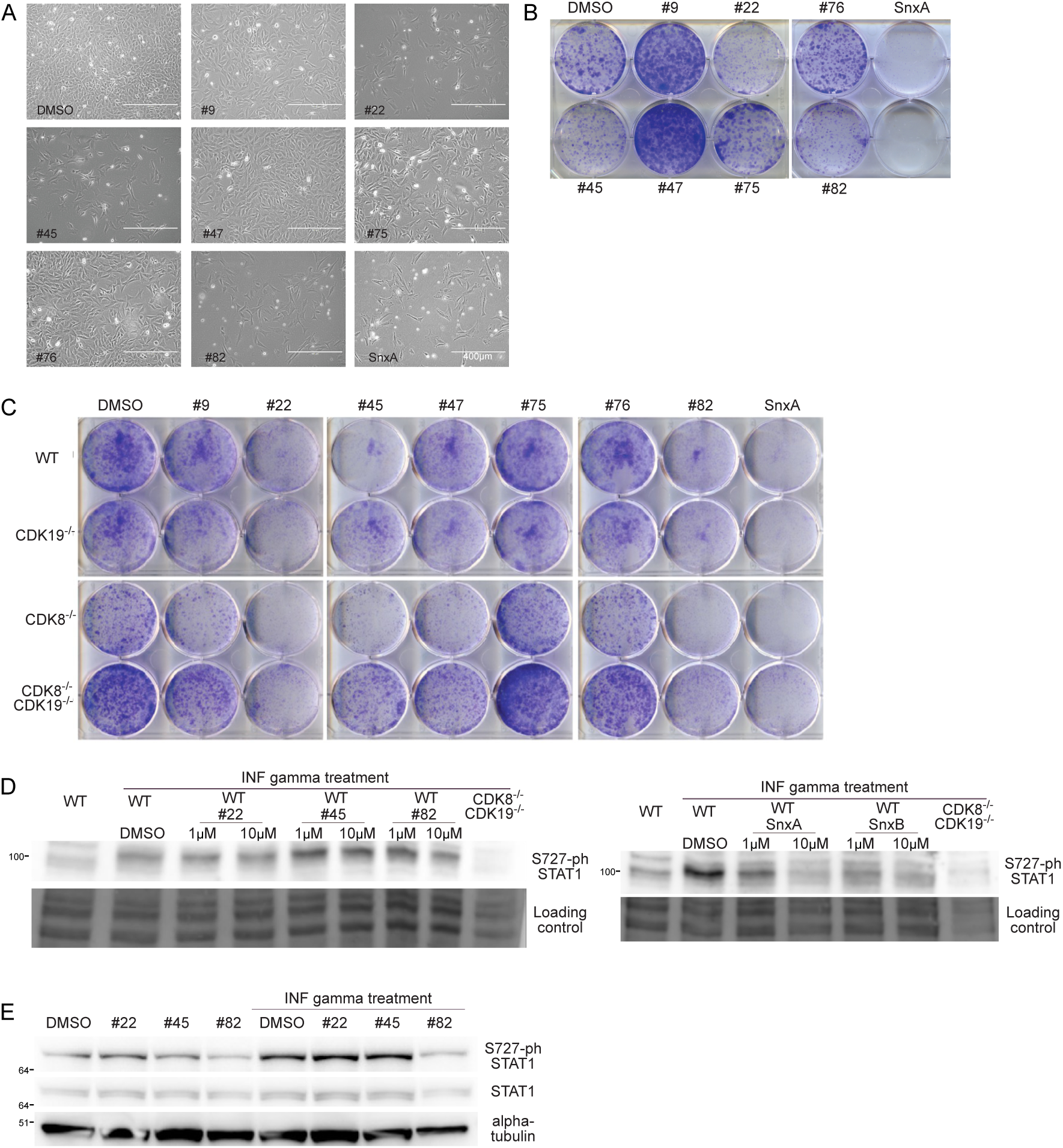
Cell toxicity and inhibition of target phosphorylation. A. MEFs seeded at 10^6^ in 10 cm plates were grown for 5 days at 10 μM of drugs and imaged. B. Colony formation assay (CFA) with WT SV-40-immortalised MEFs, seeded at 15 000 in 10cm plates, treated with inhibitors at 10 μM for 8 days. N=2. C. CFA with immortalised MEFs, WT, CDK8^-/-^, CDK19^-/-^, or double knock-out, treated with inhibitors at 10μM for 6d. N = 2. D. 3T3-immortalised MEFs were treated for 2h with the indicated inhibitors, after which γIFN (at 100 ng/ml) was added for 1h, without changing the media. Cells were collected and samples prepared for WB analysis. CDK8/19 double knockout MEF cells were used as a positive control (for the absence of STAT1-S727ph signal). N=2. E. HCT116 cells grown in spheroids were pre-incubated with γIFN (at 20 ng/ml) for 3h and treated with selected inhibitors at 10 μM or DMSO at 0.1% for 3h. Spheroids were lysed and analysed by Western blotting for the indicated proteins. Normalised STAT1-S727ph/STAT1 signal ratio are shown. N=2.

To verify whether our inhibitors effectively bind to CDK8 in the complex cellular environment, we performed cell thermal shift assays (CETSA)^61^, in which ligand binding alters the denaturation temperature and thus solubility of its target protein. We performed CETSA experiments in two different ways. In the first, MEFs were lysed and incubated with inhibitors for 15 minutes prior to denaturation, to assess whether inhibitors can access their target in a highly complex mixture. All seven inhibitors and Senexin A indeed caused an increase in the denaturation temperature of CDK8, indicating that they bind their target (Figure 5A). In the second assay, live MEFs were incubated with inhibitors for 1 hour prior to lysis and denaturation, allowing assessment of their cell permeability and target engagement in intact cells. Here, only compounds #47, #82 and Senexin A resulted in thermal stabilisation of CDK8, suggesting that other inhibitors do not effectively bind their target in living cells (Figure 5B). This corroborates cellular toxicity assays, further indicating that toxicity of compounds #22 and #45 is due to off-target effects, while lack of toxicity of #9, #75 and #76 could still in theory be due to lack of target engagement. Nevertheless, compounds #47, #82 and Senexin A show evidence for binding their target in cells. #47 presents the lowest toxicity of these, although its IC_50_ for CDK8-cyclin C is also the highest (200 nM). In conclusion, most cellular toxicity of CDK8/19 inhibitors appears to be off-target and most do not effectively bind CDK8 in living cells, while the evidence suggests that inhibition of CDK8 and CDK19 is not toxic.

**Fig. 5.**
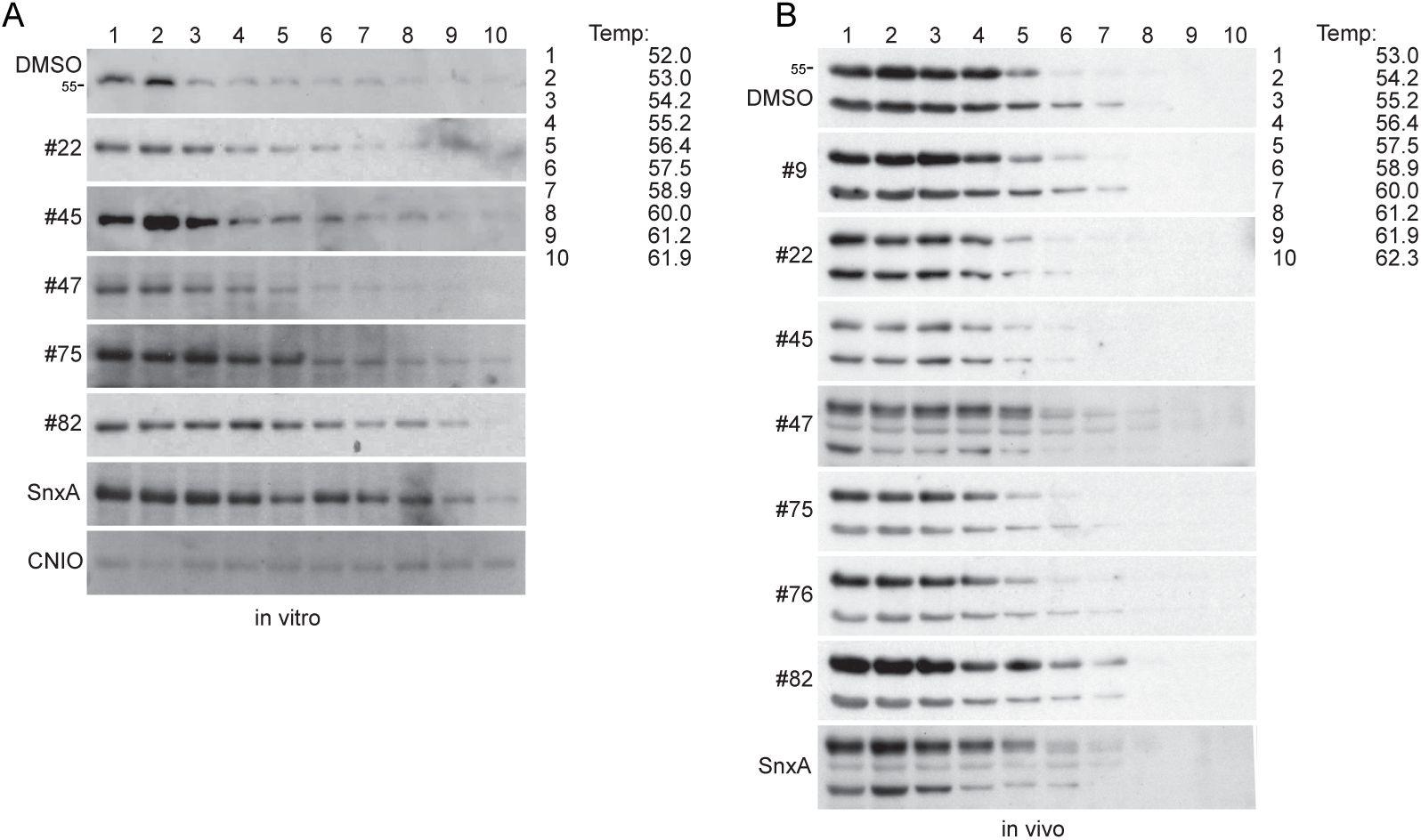
Analysis of cell penetration and target engagement of CDK8 inhibitors using CETSA assays. SV-40-immortalised MEF cell lysates (A, *in vitro* N=4) or MEF cells (B, *in vivo*, N=6) were treated with the indicated CDK8/19 inhibitors, or DMSO as a control, followed by heat treatment at the indicated temperatures and Western Blot analysis with anti-CDK8 antibody.

To further assess specificity and the reasons for biological toxicity of compounds, we used an assay that is independent of transcription: DNA replication in *Xenopus* egg extracts. In this system, transcription is silent and protein synthesis is blocked by cycloheximide treatment, while replication of DNA from added sperm chromatin is quantified by measuring incorporation of ^33^P-dCTP into insoluble material. DNA replication in *Xenopus* egg extracts is promoted by both CDK1 and CDK2 activities ^62^, but we previously observed that CDK8, CDK19 and most components of the Mediator complex are also present on chromatin ^63^. Since CDK8 was recently found to promote efficient DNA replication in mammalian cells by binding to the MTBP protein, a homologue of the essential yeast replication factor Sld7 ^37^, we assessed whether CDK8 depletion or treatment with inhibitors affects DNA replication in *Xenopus* egg extracts. We found that CDK8 binds cyclin C, as expected (Figure 6A) and is present in high molecular weight complexes (Figure 6B). We confirmed that CDK8 and cyclin C accumulate on chromatin during a 2-hour time course, in parallel with DNA replication (Figure 6C). This chromatin accumulation did not require DNA replication itself since it was not blocked by recombinant Geminin, which inhibits loading of the replicative helicase MCM components (Figure 6C). We next ascertained whether CDK8 activity is required for efficient DNA replication in this system. We removed CDK8 and cyclin C from the extract by immunoprecipitation with antibodies that cross-react with *Xenopus* CDK8 and complemented (or not) with an equivalent amount of recombinant CDK8-cyclin C protein (Figure 6D). CDK8 depletion caused a slight but non-significant reduction in rate of DNA replication when compared with mock depletion with non-specific antibodies (Figure 6E). This was not rescued by complementing with recombinant kinase, indicating that CDK8 kinase activity is not required for efficient DNA replication. To confirm this, we analysed the effect of different concentrations of Senexin A, and compared with the pan-CDK inhibitor purvalanol A. While the latter caused a 90% reduction in replication rate compared to vehicle control (Figure 6F), Senexin A had only minor effects even at high concentrations (200 µM), indicating that CDK8/CDK19 kinase activity is dispensable for efficient replication in the *Xenopus* egg extract system. We then tested compounds #9, #22, #45, #75, #76 and #82. While #9, #45 and #76 had no clear effect on DNA replication rates, #22, #75 and #82 impeded replication in a dose-dependent manner, indicating that they inhibit other activities required for DNA replication. This identifies transcription-independent off-target effects of these molecules that might contribute to toxicity. This was surprising for #75, which had low toxicity in cell models and showed no *in vitro* inhibition of CDK1, CDK2 or CDK5 activity, although the CETSA assay showed that this compound might not penetrate efficiently into cells. Thus, testing effects of kinase inhibitors on DNA replication in egg extracts is a useful method for assessing their toxicity.

**Fig. 6.**
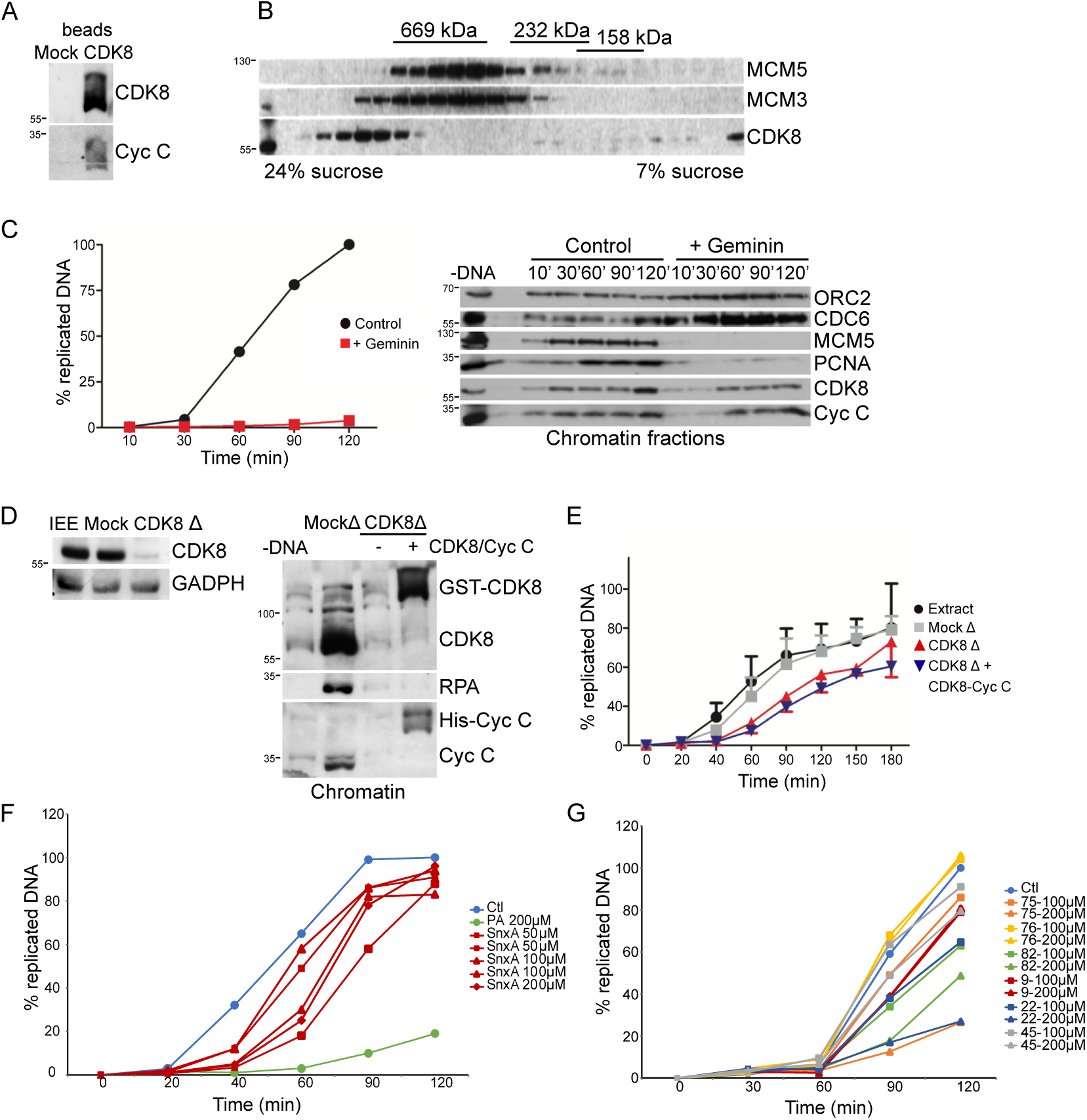
Transcriptionally silent system of Xenopus egg extract as a model for testing CDK8/19 inhibitor activity and specificity. A. CDK8 was immunoprecipitated from interphase egg extract and the beads were blotted for the indicated proteins. B. Interphase egg extract was fractionated by sucrose gradient and fractions were blotted for the indicated proteins. C. Left, chromosomal DNA replication quantified by ^32^P-dCTP incorporation assay, in control conditions, or after addition of recombinant Geminin, to block replication licencing. Right, chromatin loading of pre-replication complex (ORC2, MCM5 and CDC6) and pre-initiation complex (PCNA) replication factors, CDK8 and Cyclin C during DNA replication time course. N=4. D. Left, immunodepletion of CDK8 (but not using Mock beads) removes majority of the kinase from the interphase egg extract; GADPH was used as loading control. Right, chromatin fractions from CDK8- or Mock-immunodepleted egg extract, or CDK8-depleted extract supplemented with recombinant CDK8-Cyclin C complex were analysed by WB with the indidated proteins. Slower migrating recombinant GST-CDK8 and His-Cyclin C are indidated. E. Chromosomal DNA replication in immunodepleted extracts, as in D, was quantified. N=3. Error bars show standard deviation. F,G. Chromosomal DNA replication quantified by ^32^P-dCTP incorporation assay, in control conditions (DMSO), or in the presense of the indicated inhibitors.

To better understand the selectivity of our identified inhibitors, we reasoned that aligning CDK8-cyclin C ligand signatures for each of our compounds to all CDKs might reveal amino acids that are critical for ligand binding ^64,65^. Indeed, a single substitution of the tryptophan at position 105, which makes contacts to cortistatin A yet is not conserved among other CDKs, suffices to abrogate sensitivity of CDK8 to this compound ^21^. Figure 7 and Table S3 identify residues involved in binding of our compounds and show that most are conserved among all CDKs, suggesting that they do not confer CDK8 selectivity and might be important for catalytic activity. At the hinge region, which is a key region for binding many CDK8 inhibitors, D98, Y99 and A100 are not conserved among most CDKs, while M174, adjacent to the catalytic D173, is a phenylalanine in all other CDKs, and F176 is a leucine in all other CDKs. We incorporated substitutions of several of these residues for amino acids present in other CDKs into the structural model of CDK8-cyclin C and tested their effects on induced-fit docking of our compounds. The substitution M174F of the DMG loop had almost no effect alone, and even when combined with the adjacent F176L had only minor effects on docking of several compounds (Table S4). Hinge mutants D98E, A100L and A100M also had at best only minor effects. Surprisingly, even substitution of the residues highly conserved among all CDKs to bulky or charged residues (V35R, L70R, L158W) had negligible effects on DS of the different compounds (Table S4). Although L158R did cause a significant reduction in binding of six of the seven compounds, this would also be likely to disrupt ATP binding. This analysis suggests that the CDK8 inhibitor binding pocket is very flexible and can readily accommodate substitutions which in most cases probably do not affect kinase activity (since they are present in other CDKs) but also are not critical for binding inhibitors (Figure 7C). It also suggests that point mutations in the kinase do not readily confer resistance to our compounds, which may be important in a cancer treatment context.

**Fig. 7.**
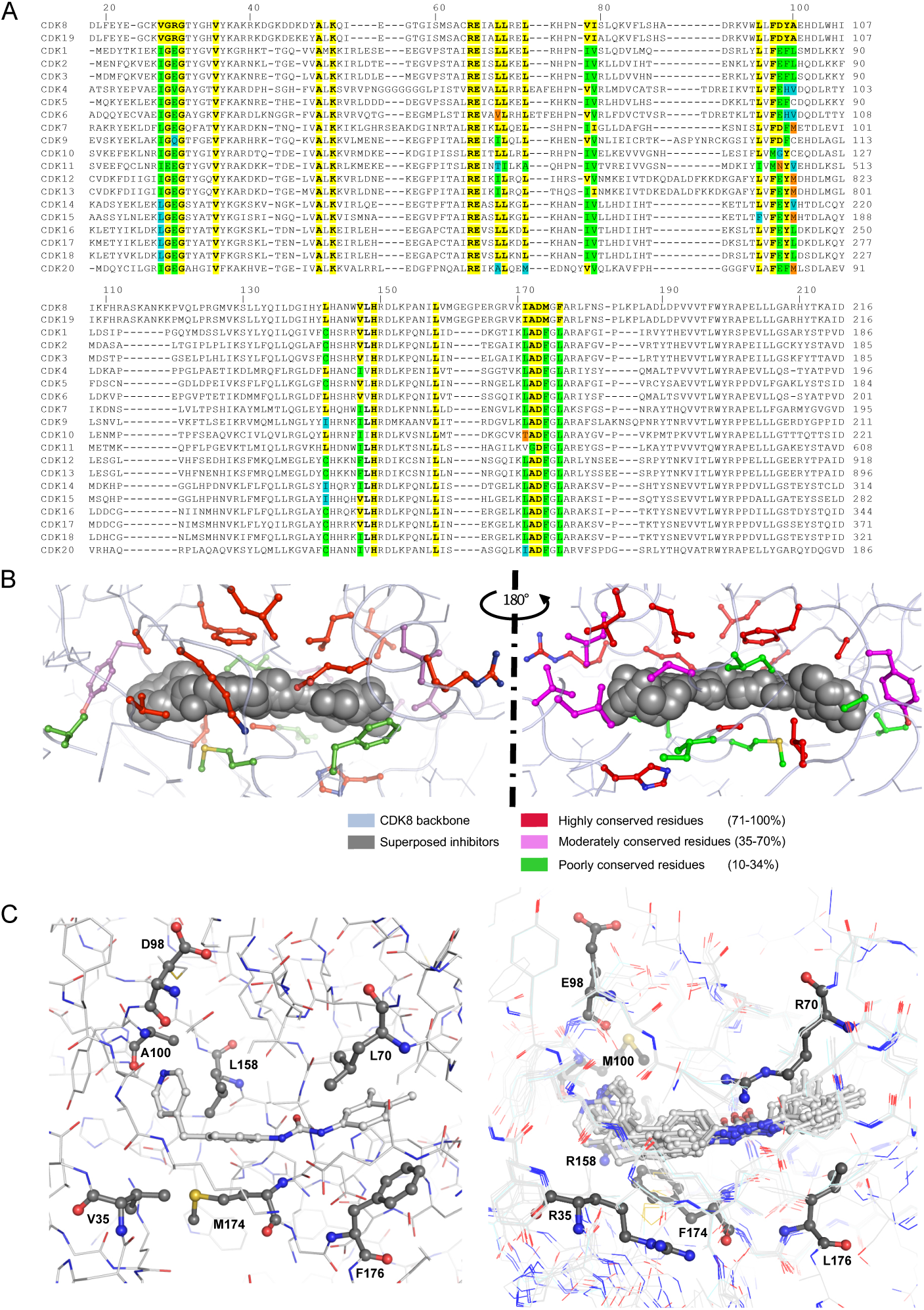
Virtual mutation of inhibitor ligand signatures reveals substantial flexibility in the kinase ATP-binding site. A. Alignment of the 20 CDK sequences. Only the regions containing the amino acids responsible for interactions with the compounds are shown. Interacting amino acids and their conservation are highlighted. B. View of the CDK8 amino acid residues involved in contact with compounds. The seven compounds are superposed and shown with gray balls. Interacting residues are depicted by colored balls and sticks according to their percentage of conservation (red: highly conserved residues (71-100%); pink: moderately conserved residues (35-70%); green: poorly conserved resides (10-34%)). C. Highlight of virtually mutated residues. Left panel: wild-type CDK8 showing the seven residues selected (ball and stick and dark grey carbon) to assess the effect of point mutations on compound binding. The induced fit pose of molecule #82 is shown (ball and stick and light grey carbon). Right panel: superposition of the poses obtained for each mutant following induced-fit docking. For clarity, only one of the two side chains is shown (ball and stick and dark grey carbon) in the case of the A100M/L and L158R/W mutants. In contrast, all the positions obtained for compound #82 are shown (ball and stick and light grey carbon).

It has recently been suggested that MED12 binding to the CDK8-cyclin C complex hinders type II inhibitor binding, probably by reorienting the DMG triplet to the “in” position ^66^, which could potentially explain the lack of binding of some of our inhibitors to CDK8 in cells. Another recent study published the structure of the entire MED12/13-containing CDK8 kinase module (CKM) along with Mediator itself ^67^. Therefore, to test whether our inhibitors lose binding affinity for the DMG-in structure and/or in the presence of MED12, we re-performed molecular docking experiments against the DMG-in structure PDB 5SX2 ^68^, and the MED12-containing CKM structure PDB 8TQ2 ^67^. We compared with the published CDK8/19 type I inhibitors cortistatin A ^21^, CCT251545 ^41^, Senexin A, B and C ^23,31,69^, as well as the non-selective type II inhibitor Sorafenib ^40^. We tested all inhibitors against both structures using both XP and induced fit parameters (Table 2 and examples shown in overlay in Figure 8A, B). As well as Sorafenib, as expected, compounds #9, #22, #45 and #47 had a docking score indicating that they bind the DMG-in structure significantly less well than the DMG-out structure used for screening, accompanied by their reorientation in the binding site. This was also the case for the type I inhibitors cortistatin A and Senexins B and C, which have established pre-clinical credentials. The DMG orientation had only minor effects on DS of #75, #76 and #82 and CCT251545 despite a similar re-orientation of the inhibitors, suggesting that these compounds have multiple modes of high affinity binding. Only Senexin A had a better DS for the DMG-in structure than for the DMG-out structure. Interestingly, all of our inhibitors and almost all published type I and type II inhibitors had significantly and similarly impaired DS against the MED12-containing CKM structure (Table 2 and Figure 8B). #82 had better a DS than cortistatin A to all three structures, supporting the notion that it is the most effective of all of our compounds and compares well with established CDK8 inhibitors, despite evidence of off-target toxicity.

**Fig. 8.**
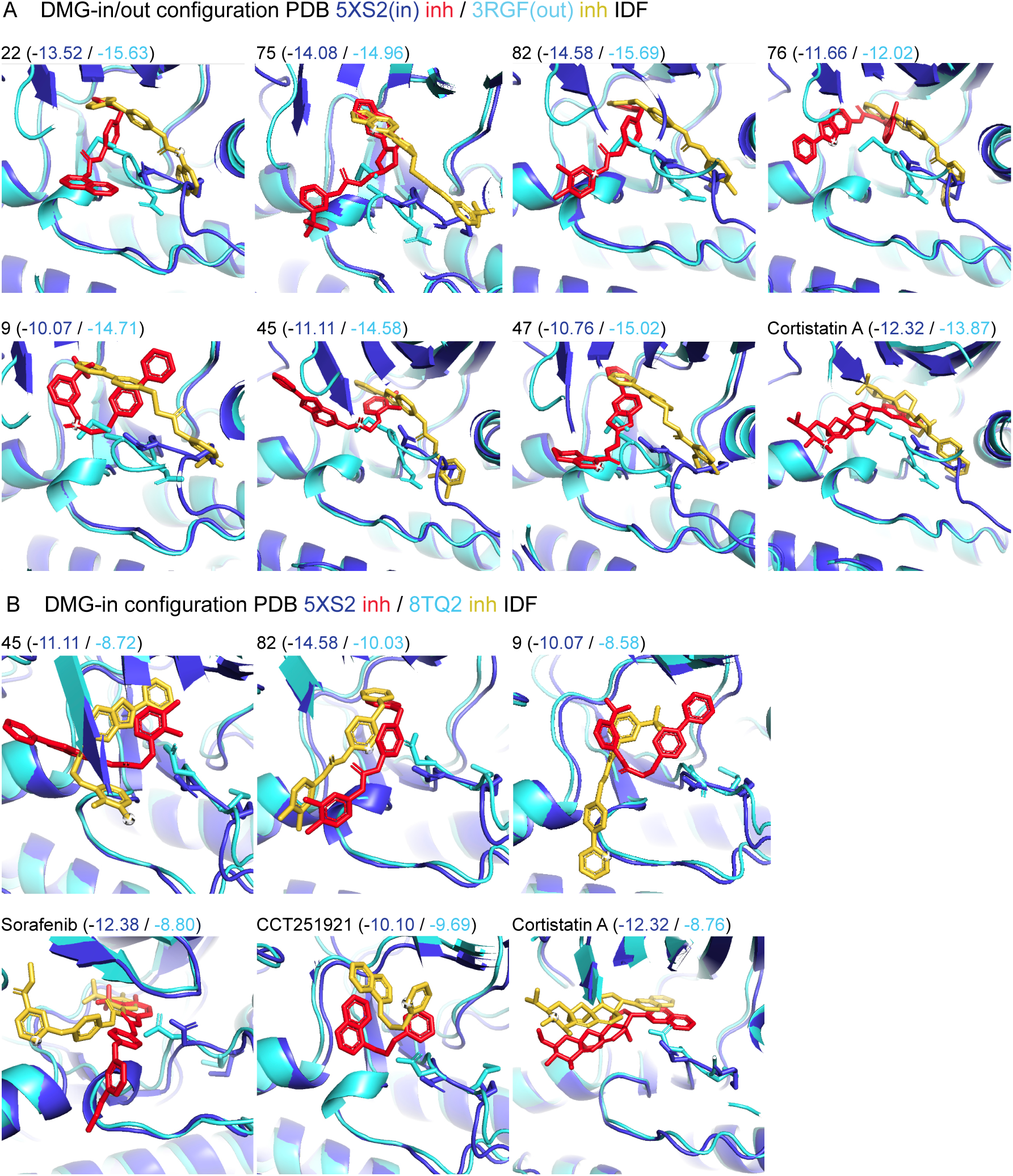
CDK8/19 inhibitors can bind CDK8 in both its inactive and active conformations. Overlay of structures of selected inhibitors with the inactive (DMG-out, PDB 3RGF; A, cyan) and active (DMG-in) conformations, without (PDB 5XS2; A and B, blue) and with MED12 (PDB 8TQ2; B, cyan). Zoom-in on ATP-binding pocket of CDK8, with side chains of D173 and M174 shown. In red, inhibitor docked into 5XS2; in yellow, inhibitor docked into 3RGF, A, and 8TQ2, B. Induced fit docking scores for compared structures are indicated.

**Table 2.**
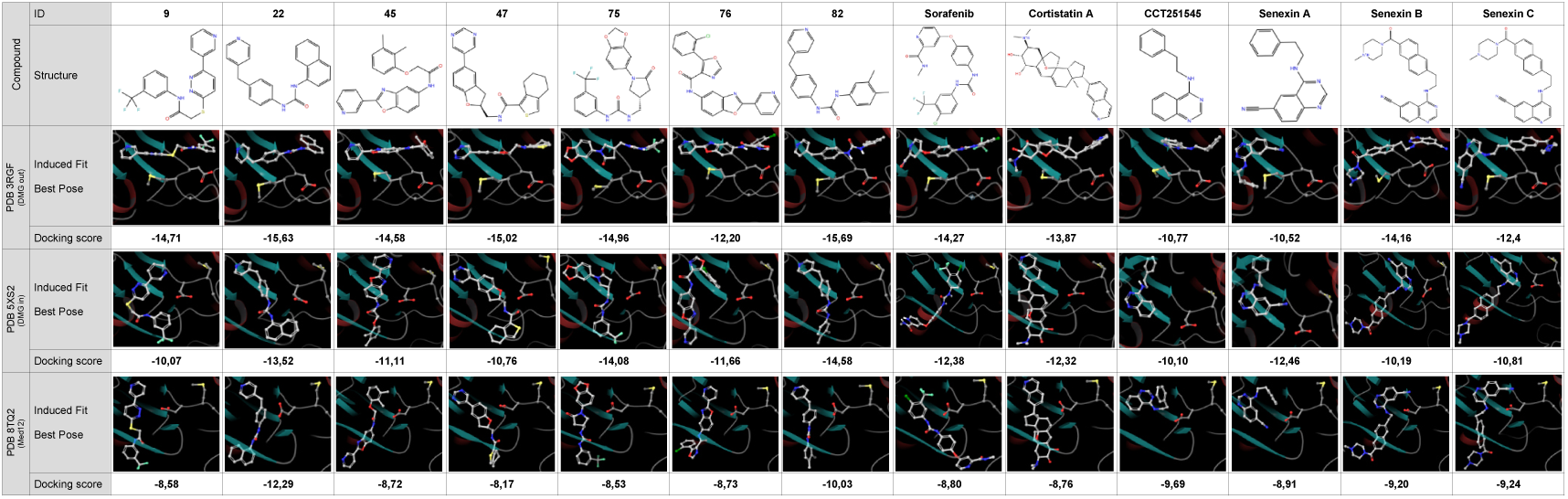
Effects of CDK8 conformations on compound binding. Induced fit docking experiments were carried out with three different CDK8 structures; a DMG-out (PDB: 3RGF), a DMG-in (PDB: 5SX2) and the MED12-containing CKM (PDB: 8TQ2). Our compounds, known CDK8 inhibitors (cortistatin A, CCT251545, Senexin A, B, C) and a non-selective inhibitor (Sorafenib) were used to assess the influence of CDK8 conformation change. Structural formula and 3D arrangement of the best docking pose for each ligand (white carbon atoms) with the CDK8 forms are shown. Amino acids of the DMG loop are represented in ball and stick.

## Discussion

We previously found that CDK8, cyclin C and Mediator complex components accumulate on chromatin during DNA replication ^63^. As it had been published that CDK8 is a colorectal cancer oncogene in Wnt-pathway dependent cancer cells, we decided to investigate their mechanisms of action and potential for CDK8 inhibitors as cancer therapeutic agents. We found that CDK8 and CDK19 are not essential for cell proliferation in the intestinal epithelium and instead modulate secretion by controlling a relatively limited subset of genes ^13^. Unexpectedly, genetic knockout of CDK8 in mice had no effect on intestinal tumourigenesis induced chemically (by AOM/DSS treatment) or genetically (by activating the Wnt pathway through *Apc* gene mutation) ^13^, and we could not confirm either the previously reported oncogenic activity ^27^ or intestinal “tumour suppressor” effect of CDK8 ^28^. Yet the presence of CDK19 might be expected to compensate for the absence of CDK8, and numerous emerging reports of CDK8/CDK19 inhibitors showed either cancer cell toxicity *in vitro* or benefit in preclinical mouse cancer models (Table 1), encouraging us to pursue development of CDK8/CDK19 inhibitors.

Indeed, CDK8 and CDK19 are considered cancer targets with high therapeutic potential. Emerging data suggest that these kinases are non-essential for most cell types, and therefore toxicity of the most specific molecules is expected to be limited, yet they appear to promote transcriptional upregulation both in amplifying oncogenic signalling ^14^ and in acquisition of therapeutic resistance to other classes of drugs. In this regard, CDK8/19 inhibitors have recently been shown to limit resistance to BCR-ABL tyrosine kinase inhibitors in models of chronic myeloid leukaemia ^70^ as well as anti-endocrine therapy or mTOR inhibitors in breast and prostate cancers ^30,32,71^ and EGFR inhibitors in several cancer models ^72^.

Here, we present a rational approach to developing new CDK8/19 inhibitors in several steps. Using molecular docking to perform virtual screening of millions of available molecules, followed by protein kinase assays of the top hits, we found that 36 out of 99 compounds indeed showed potent inhibition of CDK8-cyclin C activity *in vitro.* This validates virtual screening as a useful first approach, especially given the large-scale availability of 3D structures predicted by AlphaFold and its extended applicability to ligands ^73^. A second important finding is that the comparison of IC_50_ for inhibition of kinase activity and for cell growth is a good indicator of off-target toxicity, which was confirmed by genetic knockout of the presumed targets, an approach that is rarely performed. We also showed that the *in vitro* DNA replication system of *Xenopus* egg extracts is a simple and effective test of off-target activity in an environment close to that of living cells. Where CDK8 and its paralogue CDK19 stand on the spectrum of essentiality for cell survival and proliferation remains uncertain, but our work reported herein reinforces the emerging view that they are generally non-essential, and we could find no evidence for a direct involvement in DNA replication reported recently ^37^. We conclude that toxicity of most reported CDK8/CDK19 inhibitors is likely to be due to off-target effects, supporting conclusions of a previous study ^46^. Off-target action of drugs is far from uncommon. A recent survey used CRISPR-Cas9-mediated gene inactivation to assess 6 different apparent cancer gene dependencies targeted by 10 cancer drugs undergoing clinical trials. Cellular toxicities of all of these drugs were found to be off-targets, while the same study identified the genuine target of a putative PBK inhibitor to in fact be CDK11 ^49^. Yet since these drugs may nevertheless show clinical benefits, some toxicity may be acceptable or even desirable in order to limit resistance.

Development of potent, specific protein kinase inhibitors that have on-target activity against cancer cells with sufficiently limited toxicity remains an extremely challenging and expensive endeavour. Part of the problem is that although proteins that are directly involved in cell proliferation (e.g. the protein kinases CDK1, PLK1 and AURA) emerge as strong pan-cancer prognostic markers, no inhibitors against these targets have received regulatory approval as they are generally essential for cells and thus inhibitor toxicity is limiting; instead, approved cancer drugs tend to target non-essential proteins but as a result face frequent resistance ^74^. These fundamental issues are often compounded by insufficiently rigorous pre-clinical testing. The Reproducibility Project: Cancer Biology, only managed to replicate 50 out of 193 experiments published in high impact journals, and of these, most did not confirm the original findings ^75,76^. This suggests that pressure to find positive results and to publish in high impact journals is holding back drug development. The solution is clearly to perform more extensive fundamental research on the targets.

We present one molecule, #82, that shows comparable binding to previously discovered type I and type II inhibitors and better binding than most to the MED12-containing structure; it also has on-target activity in cells and off-target toxicity in the range of several previously reported inhibitors, despite the specificity for CDK8 compared to related kinases. Animal studies would be required to determine whether a therapeutic window is possible, but a more prudent way forward would be to explore ways of maintaining its activity while reducing toxicity.

Finally, we suggest that the dynamic flexibility of the ATP binding site renders the classification of inhibitors as type I or type II inhibitors misleading. We show evidence that not only the so-called type II inhibitor Sorafenib, but also most so-called type I inhibitors are likely to bind less well to the DMG-in conformation than the DMG-out conformation. However, the presence of MED12 in the complex more strongly affects the binding of most CDK8/19 inhibitors, and therefore such a complex should be prioritised for screening for new inhibitors.

## Materials and Methods

### Docking

Docking experiments were carried out with the Schrodinger suite (Schrodinger, LLC, New York, NY). The CDK8 / Cyclin C structures (PDB: 3RGF, 5XS2 and 8TQ2) and the CDK2 structure (PDB: 4GCJ) were used for the studies. Protein structures were processed with the Protein Preparation Wizard tool. A docking area (35Å × 35 Å × 37 Å) centered on the Sorafenib molecule from the 3RGF structure was defined to generate the docking grid. The same grid was also applied to the other CDK8 structures (PDB: 5XS2 and 8TQ2). For CDK2 (PDB: 4GCJ), an equivalent grid centered on the RC-3-89 inhibitor was defined. LigPrep was used for ligand energy minimisation and to generate ligand 3D structures, tautomers and ionisation states.

The virtual screening was performed using the ZINC 12 database and the 3RGF structure. No constraints (such as hydrogen bond or atom position) were applied to guide the binding. All compounds were docked using Glide ^57^ and the Glide docking score was used to rank the docking poses. High Throughput Virtual screening (HTVS) ^58^ was used at an early stage in order to test a large number of compounds. The best HTVS poses were then screened using the standard precision (SP) mode and finally the very best compounds were analysed with the extra precision (XP) mode ^59^.

The other docking experiments were performed using the induced fit mode. This more time-consuming mode simulates the induced conformational changes that can occur upon ligand binding providing a more accurate representation of the interactions compared to rigid docking methods. Here again, no constraints were applied and the docking score was used to rank the poses. The best of the 20 poses generated for each compound were retained.

### Cell lines

MEFs were derived from Cdk8^Lox/Lox^ Polr2a^tm1(cre/ERT2)Bbd^ and Cdk8^+/+^ Polr2a^tm1(cre/ERT2)Bbd^ embryos. Details of the creation of these mice are described ^13^. All animal work was conducted in accordance with international ethics standards and was approved by the Animal Experimentation Ethics Committee of Languedoc Roussillon. Mice were housed in a SPF animal facility according to guidelines, with no more than 5 mice per cage (501 cm^2^), which contained sawdust and wood shavings. Mice had unlimited access to food and water. MEFs were prepared according to a standard protocol (77). At E12.5, pregnant females were sacrificed and embryos collected in PBS and dissected to remove internal organs, head and limbs. The remaining carcasses were dissociated mechanically with a scalpel blade and chemically in trypsin solution; then cultured separately (P0), each embryo in a 10 cm dish. Primary cells that had adhered and proliferated in culture were amplified and then frozen. To induce the Cre-mediated recombination of *Cdk8 ^lox/lox^*, MEFs were cultured in medium supplemented with 600nM 4-Hydroxytamoxifen (Sigma, H7904) for 14 days. Evaluation of knockout efficiency was performed using genotyping, qPCR, and Western blotting. For CDK19 deletion, two days after thawing and seeding of primary MEFs *Cdk8 ^Lox/Lox^ Polr2a^tm1(cre/ERT2)Bbd^* and *Cdk8 ^+/+^ Polr2a^tm1(cre/ERT2)Bbd^*, the *Cdk19* gene was deleted by CRISPR-Cas9 as described ^77^. The *Cdk19* exon 1-targeting sgRNA sequence from Zhang’s laboratory database (5’-AAAGTGGGACGCGGCACCTA-3’) was cloned into LentiCRISPRv2 vector as described ^77^, provided by F. Zhang, Addgene plasmid #52961). Lentiviruses were produced in HEK293T cells using JetPEI (Polypus, #101-40) with empty LentiCRISPRv2 vectors (control vector, 2 μg) or sgCDK19 (2 μg), pMD2.G (1μg) and psPAX2 (1μg). The culture medium was changed 4h after transfection. HEK293T cells were incubated for 24 h; the culture medium containing the viral particles was collected and filtered through a 0,45 μm filter. Lentiviral-mediated transduction was performed twice, on two consecutive days. MEF cells were seeded in 10cm dishes and 1ml of media containing viral particles supplemented with 100 μg Polybrene (#H9268, Sigma) was added. Antibiotic selection (5 μg/ml puromycin) was maintained for 4 days. Surviving primary MEFs were then amplified and frozen. Deletion of the targeted *Cdk19* exon 1 was verified by sequencing from genomic DNA, which was extracted using KAPA Mouse genotyping kit (Clinisciences, KK7352).

MEFs were cultured under hypoxic conditions at 1% O2 and 5% CO2. Other cells were grown at 21% O2 and 5% CO_2_ in humidified conditions. HEK293T (source, ATCC, # CRL-3216), SH-SY5Y (source, ECACC, #94030304) and U2OS cells (source, ATCC, #HTB-96) were cultured in DMEM culture medium (Dulbecco’s Modified Eagle Medium, High glucose, GlutaMAX, 31966047, Life Technologies) + 10% fetal calf serum (D7524, Sigma or Gibco) + 100 IU/ml-100 μg/ml Penicillin-Streptomycin. HCT116 cells (source, ATCC, #CCL-247) were cultured in McCoy’s medium (Life Technologies) + 10% fetal calf serum; hTERT RPE-1 cells (source, ATCC, #CRL-4000) in DMEM/F-12 medium (Life Technologies) + 10% fetal calf serum.

Mycoplasma testing was performed weekly.

### Western blotting

Protein samples were suspended in Laemmli buffer, run on SDS-PAGE gels, transferred to nitrocellulose, blocked with 3% bovine serum albumin in TBS-0.1% Tween-20. Immunoblotting was performed using unlabeled primary antibodies for 1-4 hours in the same buffer at room temperature, and horseradish-peroxidase conjugated species-specific anti IgG antibodies for one hour. Staining used enhanced chemiluminescence reagents from Perkin-Elmer, and exposures were captured on an Amersham Imager 680.

### Cell viability assessment

Cells were grown in 96-well plates in the presence of increasing concentrations of each compound (from 0.05 to 50 µM) for 48 h. Cell viability was then assessed using the CellTiter96 AQueous cell proliferation assay from Promega, according to manufacturer’s instructions. Each experiment was performed in triplicate and EC_50_ were determined from the dose-response curves according to the signal given by the control set at 100% viability, using GraphPad Prism software.

### Spheroids

HCT116 spheroids were produced using a liquid overlay technique as described ^78^. Briefly, 96-well flat-bottom plates were coated with 60 μl of agarose-DMEM (1.5% wt/vol), then extemporaneously loaded with 200 μl/well of HCT116 at 18,000 cells/ml concentration. Following a 4-day incubation period, until spheroids reached an average 400 µm in diameter, spheroids were pre-incubated with γIFN at 20 ng/ml for 3h, followed by an incubation with the compounds for 3h at 10 μM (0.1% DMSO). Spheroids were then lysed on ice in Lysis Buffer (Tris-HCl pH7.5 at 50 mM, NaCl at 300 mM, 1% NP40, β-glycerophosphate at 20 mM, Na-vanadate at 0.3 mM, Na-pyrophosphate at 5 mM, Na-fluoride at 5 mM, PMSF 0.5 mM, 0.5% sodium deoxycholate, benzonase at 40 U/μl and protease inhibitor cocktail), sonicated and centrifuged. Protein concentration in clear lysates was determined by Bradford assay before being subjected to SDS-PAGE electrophoresis and transfer on nitrocellulose.

### MEF immortalisation

3T3 method. MEFs were passaged multiple times, transfering 3x10^5^ cells every third day into a new plate ^79^. SV-40 method. 1.5 x10^6^ HEK293T cells were seeded in 10 cm plates. The following day, when cells reached 70% confluency, cells were transfected with a mix containing the retroviral vector pBABE-puro SV40 LT (containing SV40 simian virus T40 antigen; Addgene), pFB Moloney plasmid (containing ecotropic envelop; Agilent), pGag pol plasmid (containing polymerase; Addgene) and jetPEI (Polyplus) transfection reagent, according to the Polyplus protocol. 36h and 48h after transfection, viruses were collected from the media, filtered through a 45 μm filter and frozen at - 80°C. MEFs were plated at 8x10^5^ cells/plate (10 cm plate). 5 hours later, 6 ml of virus-containing media and 4 μg/ml of polybrene were added on top of the MEFs. A second round of infection was performed the next day. 24h later, virus-containing media were replaced with fresh media. Puromycin selection (1.5 ng/μl) was performed for 2 days.

### CETSA assays

CETSA experiments *were* performed according to Jafari et al. ^80^. For *in vitro* assays, SV-40-immortalised MEF cells were pelleted, whole cell extract prepared, drugs were added at 200μM and incubated for 15min. Heat treatment was performed for 5min in the heating block. For *in vivo* assays, SV-40-immortalised MEFs were incubated with inhibitors at 20μM for 1h. Heat treatment was performed for 3 min, followed by whole cell extraction for WB analysis.

### Colony formation assay (CFA)

SV-40-immortalised MEFs were seeded at 15 000 in 10cm plates, the inhibitors were added on the following day at 10 µM. 8 (Figure 3B) or 6 (Figure 3C) days later, cells were fixed with 4% formaldehyde, stained with crystal violet (0.5% w/v) and imaged at 10-fold magnification.

### Xenopus egg extracts

Preparation of *Xenopus laevis* egg extracts and chromatin fractions, depletions, DNA replication assays are described ^62^. Cdk8 was immunodepleted from egg extracts using protein A Dynabeads (Life Technology) saturated with anti-Cdk8 antibody or pre-immune rabbit IgG (mock depletion). Two rounds of 20 minutes depletion were performed at 4°C. Recombinant Geminin (gift from M. Lutzmann, IGH Montpellier) was used at 80 nM final concentration. Recombinant CDK8 – Cyclin C complex (ProQinase, # 0376-0390-1) was used at 100 dilution, 8,36 ng/ul final. Sucrose gradient. 100 μl of interphase egg extract was diluted in 100 μl of XB buffer (100 mM KCl, 0.1 mM CaCl_2_, 1 mM MgCl_2_, 10 mM potassium HEPES pH 7.7 (with KOH), 50 mM sucrose, 1mM DTT) + protease inhibitors (cocktail Sigma), layered over sucrose gradient 24-7% and centrifuged for 22h, at 26 000 rpm at 4°C (in SWI55TI rotor). 200 μl fractions were collected from bottom to top and analysed by WB.

### Active CDK8/Cyclin C expression and purification

Human CDK8 with N-terminal His 8x-tag and human Cyclin C N-terminally fused to GST-TEV-3C protease cleavage sites were cloned into pFastBac Dual (Invitrogen). Recombinant bacmid and baculovirus were generated and both proteins were coexpressed in Sf9 insect cells and purified on an Ni-HiTrap column by GenScript.

### Protein kinase assays

Kinase assays were performed in various buffers (see below), according to the enzyme specificity, in the presence of 15 µM [γ-^33^P] ATP (3,000 Ci/mmol; 10 mCi/ml) for 30 min at 30°C. Controls were performed with appropriate dilutions of dimethylsulfoxide. Kinase details: *Hs*CDK8/Cyclin C (expressed in Sf9 insect cells) was assayed in buffer J with 0.35 μg/μl of RBER-IRStide (STRPPTLSPIPHIPRSPYKFPSSPLRIPGGNIYISPLKSPYKISEGLPTPTKMTPRSRILVSIGESFGTSEKFQKINQMVCNSDRVLKRSAEGSNPPKPLKKLRFDIEGSDEADGSKHLPGESKFQQKLAEMTSTRTRMQKQKMNDSMDTSNKEEKRRRRRRRRRRRHTDDGYMPMSPGVA, inspired from ProQinase, synthesized by GeneCust, expressed in bacteria) as substrate; *<u>Hs</u>*<u>PIM1</u> (expressed in bacteria) was assayed in buffer B with 0.8 μg/μl of histone H1 (Sigma #H5505) as substrate. *<u>Hs</u>*<u>Haspin-kd</u> (kinase domain, amino acids 470 to 798, expressed in bacteria) was assayed in buffer H with 0.007 μg/μl of Histone H3 (1-21) peptide (ARTKQTARKSTGGKAPRKQLA) as substrate; *<u>Hs</u>*<u>CDK2/CyclinA</u> (cyclin-dependent kinase-2, human, kindly provided by Dr. A. Echalier-Glazer, Leicester, UK) was assayed in buffer A (+ 0.15 μg/μl of BSA + 0.23 μg/μl of DTT) with 0.8 μg/μl of histone H1 as substrate. *<u>Hs</u>*<u>CDK5/p25</u> (expressed in bacteria) was assayed in buffer B, with 0.8 μg/μl of histone H1 as substrate. *<u>Ssc</u>*<u>GSK-</u> <u>3</u>α/β (native glycogen synthase kinase-3 from porcine brain, affinity purified) was assayed in buffer A (+ 0.15 μg/μl of BSA + 0.23 μg/μl of DTT), with 0.010 μg/μl of GS-1 peptide, a GSK-3-selective substrate (YRRAAVPPSPSLSRHSSPHQSpEDEEE). *<u>Rn</u>*<u>DYRK1A-kd</u> (*Rattus norvegicus,* amino acids 1 to 499 including the kinase domain, expressed in bacteria, DNA vector kindly provided by Dr. W. Becker, Aachen, Germany) was assayed in buffer A (+ 0.5 μg/μl of BSA + 0.23 μg/μl of DTT) with 0.033 μg/μl of the following peptide: KKISGRLSPIMTEQ as substrate; *<u>Mm</u>*<u>CLK1</u> (from *Mus musculus*, expressed in bacteria) was assayed in buffer A (+ 0.15 μg/μl of BSA + 0.23 μg/μl of DTT) with 0.027 μg/μl of the following peptide: GRSRSRSRSRSR. Peptide substrates were obtained from Proteogenix (Oberhausbergen, France). <u>Buffer details:</u> (A) 10 mM MgCl_2_, 1 mM EGTA, 1 mM DTT, 25 mM Tris-HCl pH 7.5, 50 µg/mL heparin; (B) 60 mM β-glycerophosphate, 30 mM p-nitrophenyl-phosphate, 25 mM MOPS (pH 7), 5 mM EGTA, 15 mM MgCl_2_, 1 mM DTT, 0.1 mM sodium orthovanadate; (D) 25 mM MOPS, pH7.2, 12.5 mM β-glycerophosphate, 25 mM MgCl_2_, 5 mM EGTA, 2 mM EDTA, 0.25 mM DTT; (H) MOPS 25 mM pH 7.5, 10 mM MgCl_2_; (J) HEPES 50 mM, MgCl_2_ 10 mM, MnCl_2_ 3 mM, EGTA 1mM, DTT 2 mM, Tween 0.01 %.

### Antibodies

CDK8 - Santa Cruz, sc-13155

CDK19 – Sigma, HPA007053

Cyclin C – Abcam, ab85927

STAT1 S727-ph - Cell Signaling, #9177

STAT1-Cell Signaling, #9172

Alpha-Tubulin (clone B-5-1-2), Sigma #T5168

XMcm3, XOrc2, XRPA, XCdc6 – gifts from M. Méchali

MCM5 – Abcam, CRCT5.1 (A2.7A3), ab6154

PCNA – Abcam, ab18197

### Reagents

IFN gamma – Invitrogen, BMS326

Senexin A and Senexin B were gifts from Igor Roninson.

## Data availability statement

The datasets generated during and/or analysed during the current study are available from the corresponding author on reasonable request.

## Acknowledgements

This work was funded by the French National Cancer Institute (INCa, grants PLBIO10-068 and PLBIO15-005), and the Ligue contre le Cancer (EL2010.LNCC/DF, EL2013.LNCC/DF and EL2018.LNCC/DF). It was granted access to the CCRT High-Performance Computing (HPC) facility under the Grant CCRT2025-borelf awarded by the Fundamental Research Division (DRF) of CEA. Thanks to Pierre Brindeau for his help in drawing compound structures.

---

**Table S1.**
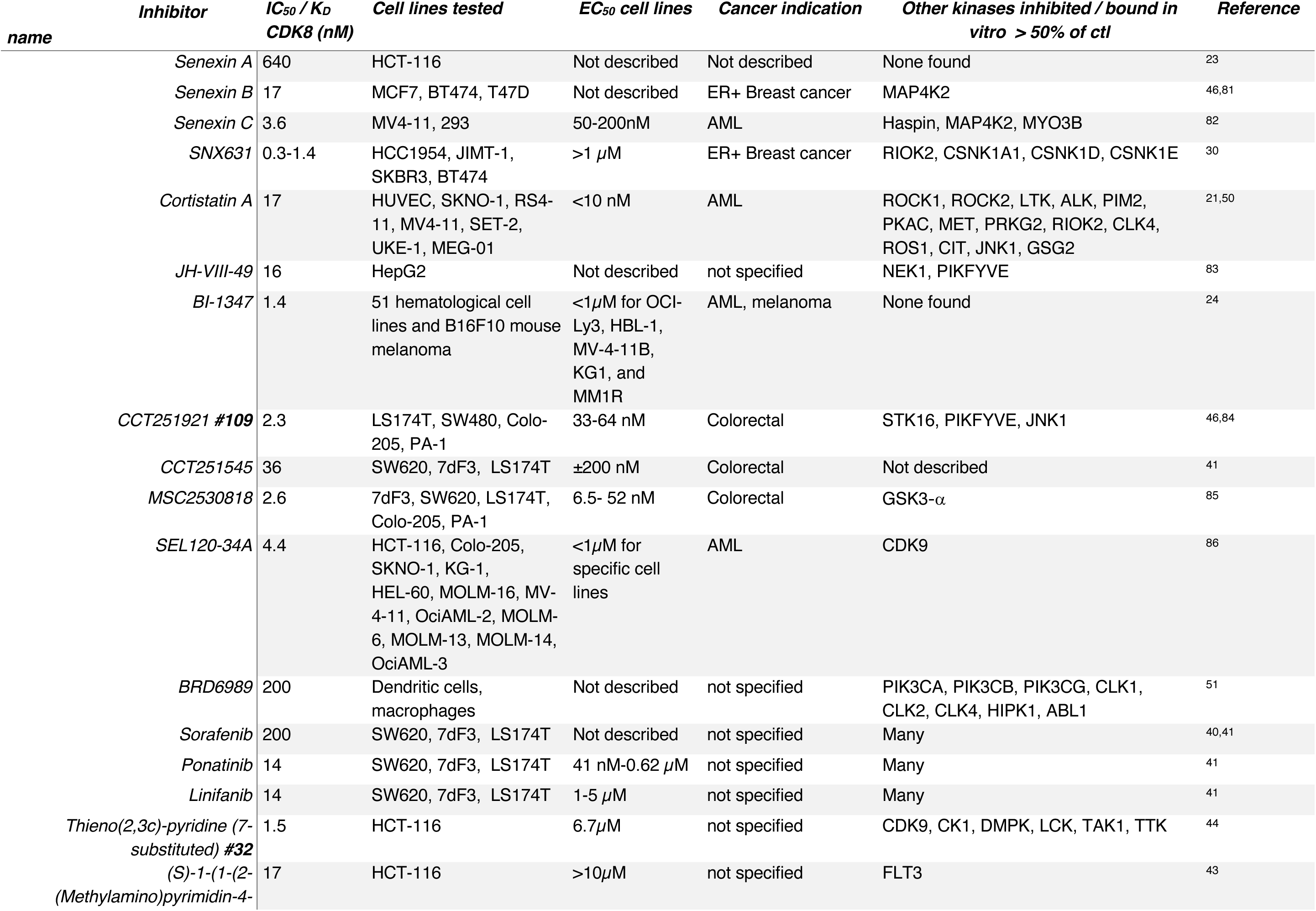
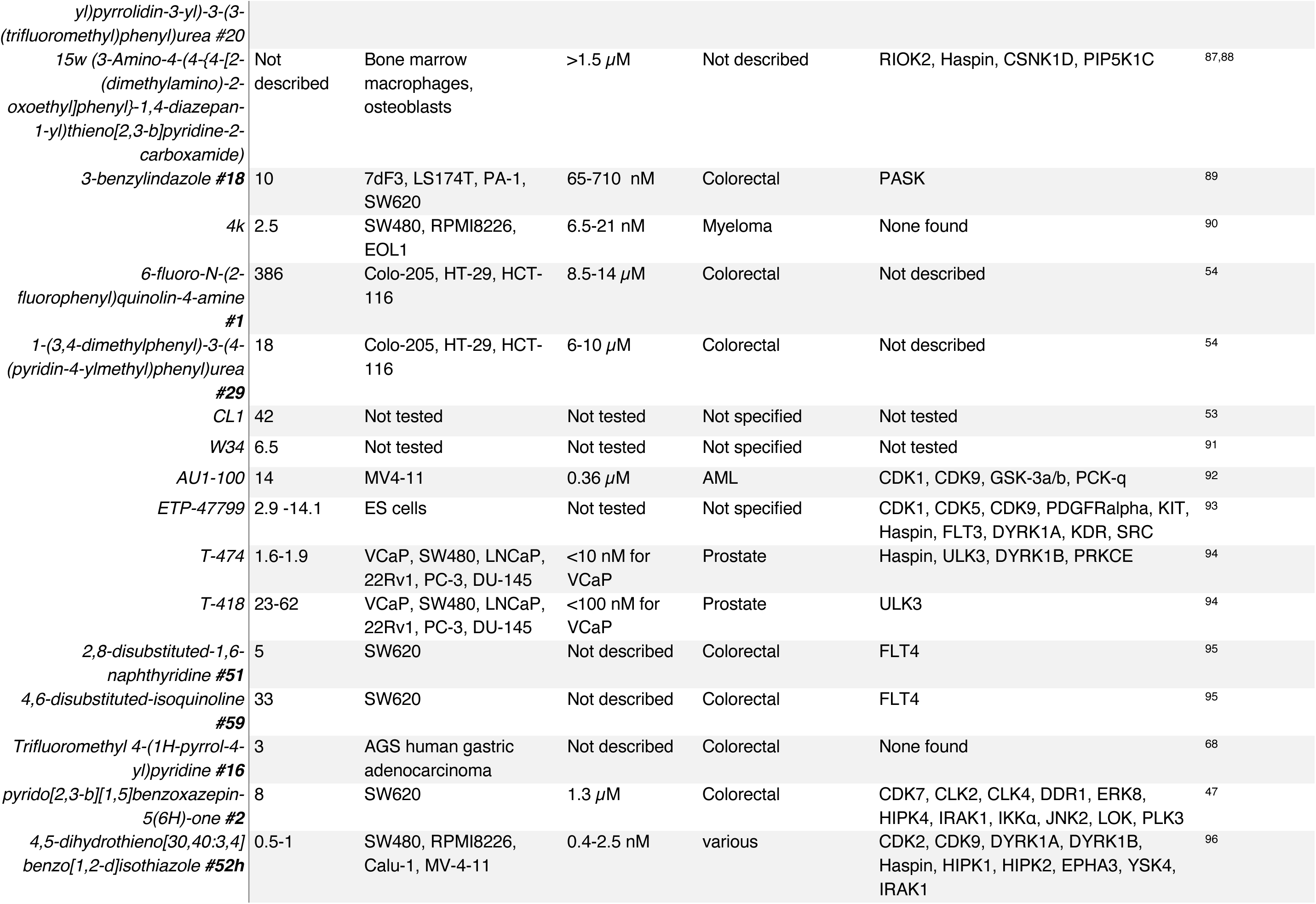
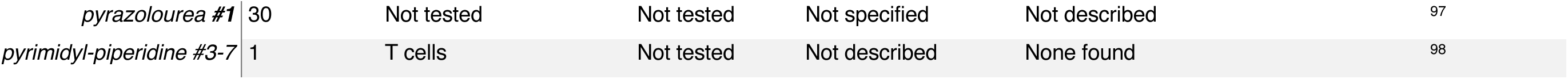
Summary of CDK8/19 inhibitors published to-date. A single molecule with the best characteristics from each chemical series is chosen from each publication.

| Table S1<br>name | Inhibitor | IC<sub>50</sub> / K<sub>D</sub><br>CDK8 (nM) | Cell lines tested | EC<sub>50</sub> cell lines | Cancer indication | Other kinases inhibited / bound in<br>vitro > 50% of ctrl | Reference |
| --- | --- | --- | --- | --- | --- | --- | --- |
|  | Senexin A | 640 | HCT-116 | Not described | Not described | None found | 23 |
|  | Senexin B | 17 | MCF7, BT474, T47D | Not described | ER+ Breast cancer | MAP4K2 | 46,81 |
|  | Senexin C | 3.6 | MV4-11, 293 | 50-200nM | AML | Haspin, MAP4K2, MYO3B | 82 |
| | SNX631 | 0.3-1.4 | HCC1954, JIMT-1,<br>SKBR3, BT474 | >1 μM | ER+ Breast cancer | RIOK2, CSNK1A1, CSNK1D, CSNK1E | 30 |
|  | Cortistatin A | 17 | HUVEC, SKNO-1, RS4-11, MV4-11, SET-2,<br>UKE-1, MEG-01 | <10 nM | AML | ROCK1, ROCK2, LTK, ALK, PIM2,<br>PKAC, MET, PRKG2, RIOK2, CLK4,<br>ROS1, CIT, JNK1, GSG2 | 21,50 |
|  | JH-VIII-49 | 16 | HepG2 | Not described | not specified | NEK1, PIKFYVE | 83 |
| | BI-1347 | 1.4 | 51 hematological cell<br>lines and B16F10 mouse<br>melanoma | <1μM for OCI-Ly3, HBL-1,<br>MV-4-11B,<br>KG1, and<br>MM1R | AML, melanoma | None found | 24 |
|  | CCT251921 #109 | 2.3 | LS174T, SW480, Colo-205, PA-1 | 33-64 nM | Colorectal | STK16, PIKFYVE, JNK1 | 46,84 |
| | CCT251545 | 36 | SW620, 7dF3, LS174T | ±200 nM | Colorectal | Not described | 41 |
| | MSC2530818 | 2.6 | 7dF3, SW620, LS174T,<br>Colo-205, PA-1 | 6.5- 52 nM | Colorectal | GSK3-α | 85 |
| | SEL120-34A | 4.4 | HCT-116, Colo-205,<br>SKNO-1, KG-1,<br>HEL-60, MOLM-16, MV-4-11, OciAML-2, MOLM-6, MOLM-13, MOLM-14,<br>OciAML-3 | <1μM for<br>specific cell<br>lines | AML | CDK9 | 86 |
|  | BRD6989 | 200 | Dendritic cells,<br>macrophages | Not described | not specified | PIK3CA, PIK3CB, PIK3CG, CLK1,<br>CLK2, CLK4, HIPK1, ABL1 | 51 |
|  | Sorafenib | 200 | SW620, 7dF3, LS174T | Not described | not specified | Many | 40,41 |
| | Ponatinib | 14 | SW620, 7dF3, LS174T | 41 nM-0.62 μM | not specified | Many | 41 |
| | Linifanib | 14 | SW620, 7dF3, LS174T | 1-5 μM | not specified | Many | 41 |
| | Thieno(2,3c)-pyridine (7-substituted) #32 | 1.5 | HCT-116 | 6.7μM | not specified | CDK9, CK1, DMPK, LCK, TAK1, TTK | 44 |
| | (S)-1-(1-(2-<br>(Methylamino)pyrimidin-4- | 17 | HCT-116 | >10μM | not specified | FLT3 | 43 |

Table S1. *Summary of CDK8/19 inhibitors published to-date.*
|  |  |  |  |  |  |  |
| --- | --- | --- | --- | --- | --- | --- |
| yl)pyrrolidin-3-yl)-3-(3-(trifluoromethyl)phenyl)urea #20 |  |  |  |  |  |  |
| 15w (3-Amino-4-(4-{4-[2-(dimethylamino)-2-oxoethyl]phenyl}-1,4-diazepan-1-yl)thieno[2,3-b]pyridine-2-carboxamide) | Not described | Bone marrow macrophages, osteoblasts | >1.5 μM | Not described | RIOK2, Haspin, CSNK1D, PIP5K1C | 87,88 |
| 3-benzylindazole #18 | 10 | 7dF3, LS174T, PA-1, SW620 | 65-710 nM | Colorectal | PASK | 89 |
| 4k | 2.5 | SW480, RPMI8226, EOL1 | 6.5-21 nM | Myeloma | None found | 90 |
| 6-fluoro-N-(2-fluorophenyl)quinolin-4-amine #1 | 386 | Colo-205, HT-29, HCT-116 | 8.5-14 μM | Colorectal | Not described | 54 |
| 1-(3,4-dimethylphenyl)-3-(4-(pyridin-4-ylmethyl)phenyl)urea #29 | 18 | Colo-205, HT-29, HCT-116 | 6-10 μM | Colorectal | Not described | 54 |
| CL1 | 42 | Not tested | Not tested | Not specified | Not tested | 53 |
| W34 | 6.5 | Not tested | Not tested | Not specified | Not tested | 91 |
| AU1-100 | 14 | MV4-11 | 0.36 μM | AML | CDK1, CDK9, GSK-3a/b, PCK-q | 92 |
| ETP-47799 | 2.9 -14.1 | ES cells | Not tested | Not specified | CDK1, CDK5, CDK9, PDGFRalpha, KIT, Haspin, FLT3, DYRK1A, KDR, SRC | 93 |
| T-474 | 1.6-1.9 | VCaP, SW480, LNCaP, 22Rv1, PC-3, DU-145 | <10 nM for VCaP | Prostate | Haspin, ULK3, DYRK1B, PRKCE | 94 |
| T-418 | 23-62 | VCaP, SW480, LNCaP, 22Rv1, PC-3, DU-145 | <100 nM for VCaP | Prostate | ULK3 | 94 |
| 2,8-disubstituted-1,6-naphthyridine #51 | 5 | SW620 | Not described | Colorectal | FLT4 | 95 |
| 4,6-disubstituted-isoquinoline #59 | 33 | SW620 | Not described | Colorectal | FLT4 | 95 |
| Trifluoromethyl 4-(1H-pyrrol-4-yl)pyridine #16 | 3 | AGS human gastric adenocarcinoma | Not described | Colorectal | None found | 68 |
| pyrido[2,3-b][1,5]benzoxazepin-5(6H)-one #2 | 8 | SW620 | 1.3 μM | Colorectal | CDK7, CLK2, CLK4, DDR1, ERK8, HIPK4, IRAK1, IKKα, JNK2, LOK, PLK3 | 47 |
| 4,5-dihydrothieno[30,40:3,4]benzo[1,2-d]isothiazole #52h | 0.5-1 | SW480, RPMI8226, Calu-1, MV-4-11 | 0.4-2.5 nM | various | CDK2, CDK9, DYRK1A, DYRK1B, Haspin, HIPK1, HIPK2, EPHA3, YSK4, IRAK1 | 96 |
| *pyrazolourea #1* | 30 | Not tested | Not tested | Not specified | Not described | 97 |
| *pyrimidyl-piperidine #3-7* | 1 | T cells | Not tested | Not described | None found | 98 |

**Table S2.**
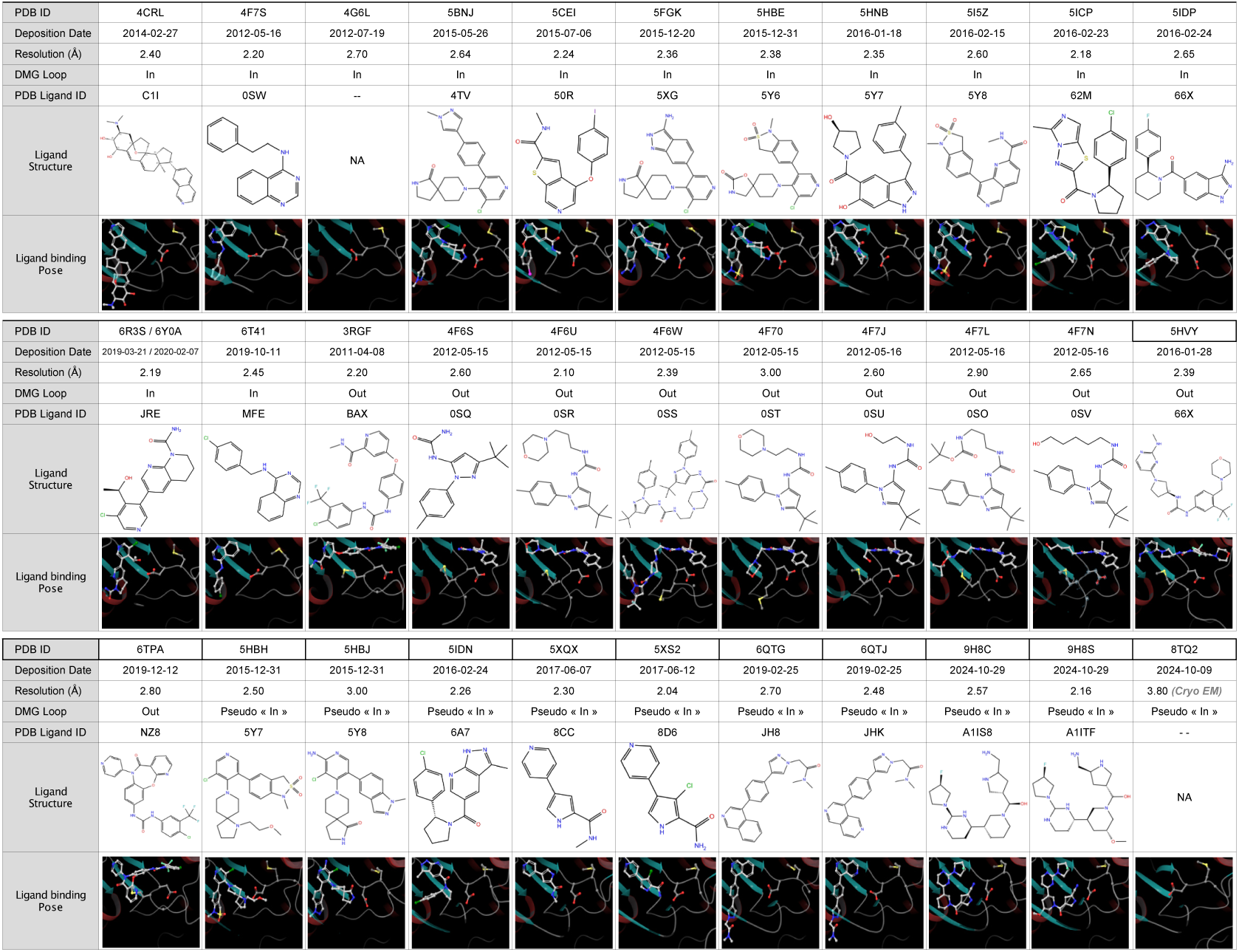
Available structures of CDK8 with inhibitors. Overview of CDK8 structures deposited in the PDB database. The structural formula and 3D arrangement of the ligands (white carbon atoms) into CDK8 active site are shown. The conformation of the DMG loop is indicated and the amino acids that form it are highlighted (ball and stick). Both “in” and “out” conformations have been described for the DMG loop of CDK8, but some structures exhibit a loop position that is not “out” and not really “in”; these structures have been classified as “pseudo in”.

**Table S3.**
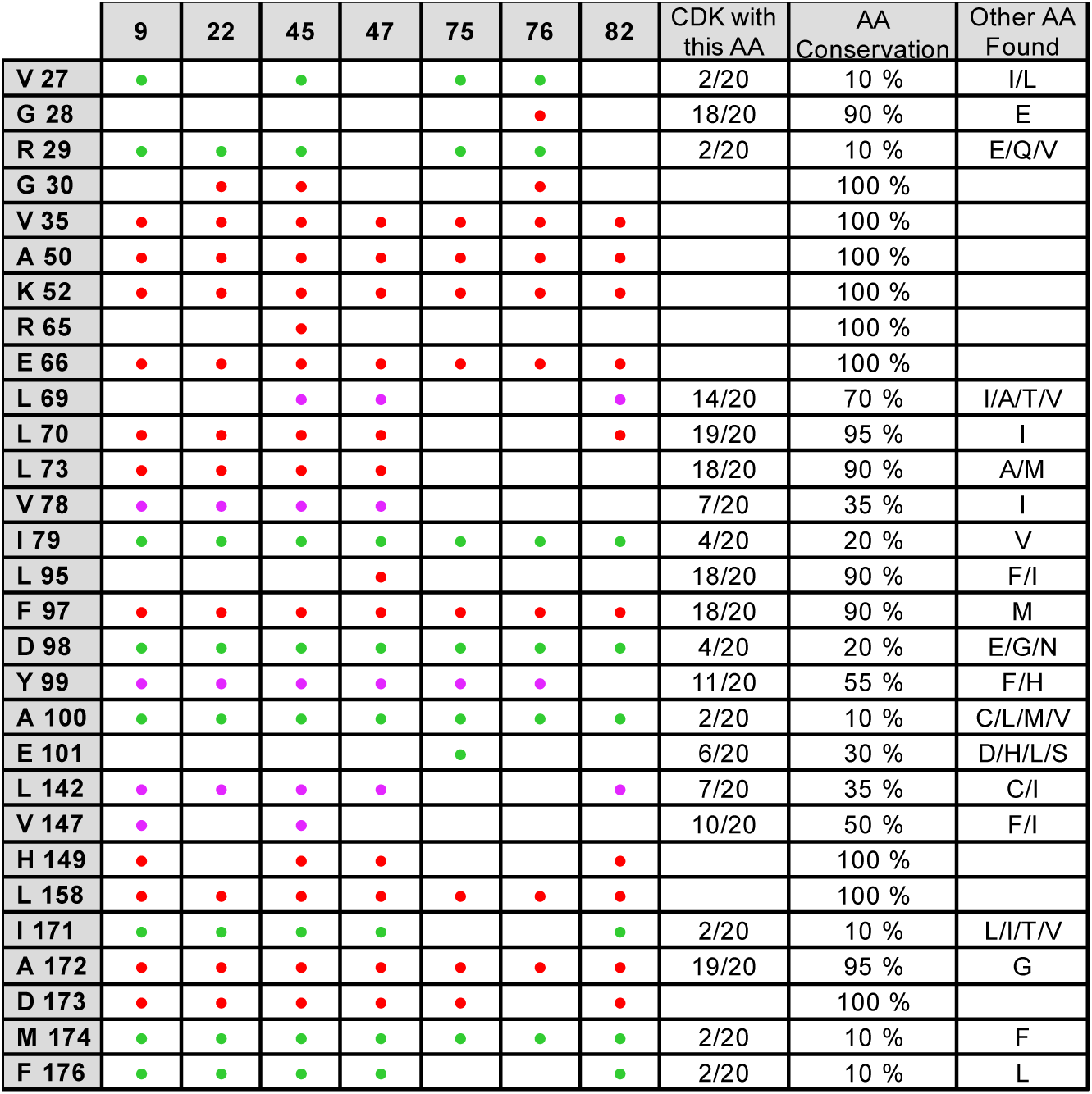
Table summarising the ligand signatures of our compounds. Contacts established by each amino acid with each compound and the percentage of conservation of each of these among all CDKs (red: highly conserved residues (71-100%); pink: moderately conserved residues (35-70%); green: poorly conserved resides (10-34%)).

**Table S4.** Effects of CDK8 substitutions on inhibitor docking. Two types of mutations were virtually generated. The “consensus” mutations(1), which replace a non-conserved residue of CDK8 with a residue conserved among other CDKs, and the “steric” mutations(2), designed to obstruct the binding site and thus prevent compound binding. Induced fit docking experiments were carried out using CDK8 (PDB: 3RGF) structure as protein target to evaluate the effect of these mutations. Docking scores corresponding to the best docking poses are reported.

| Compound | ZINC | WT | V35R | D98E | A100L | A100M | L158R | L158W | M174F | A100M<br>A155Q<br>M174F | A155Q<br>M174F | M174F<br>F176L |
| --- | --- | --- | --- | --- | --- | --- | --- | --- | --- | --- | --- | --- |
| 82 | ZINC00250053 | -15,22 | -13,55 | -15,14 | -13,54 | -15,11 | -12,09 | -15,2 | -14,37 | -15,26 | -15,04 | -13,78 |
| 22 | ZINC01197119 | -15,65 | -14,43 | -15,31 | -13,77 | -15,2 | -12,56 | -15,19 | -15,35 | -16,34 | -15,26 | -13,88 |
| 45 | ZINC02375550 | -14,44 | -14,34 | -14,07 | -13,04 | -14,78 | -13,4 | -12,3 | -14,18 | -12,64 | -14,24 | -14,11 |
| 9 | ZINC04391911 | -14,52 | -13,61 | -14,8 | -14,82 | -14,79 | -13,46 | -15,08 | -14,17 | -14,81 | -14,74 | -13,52 |
| 75 | ZINC09065754 | -14,83 | -14,30 | -15,76 | -14,79 | -15,12 | -14,52 | -15,94 | -15,11 | -15,1 | -15,5 | -14,5 |
| 47 | ZINC12040804 | -15,11 | -15,24 | -14,98 | -15,41 | -15,29 | -13,71 | -15,86 | -14,66 | -15,69 | -14,59 | -14,01 |
| 76 | ZINC19368291 | -14,40 | -12,43 | -13,95 | -13,99 | -12,11 | -13,8 | -12,68 | -14,35 | -12,74 | -14,06 | -12,92 |

**Fig. S1.**
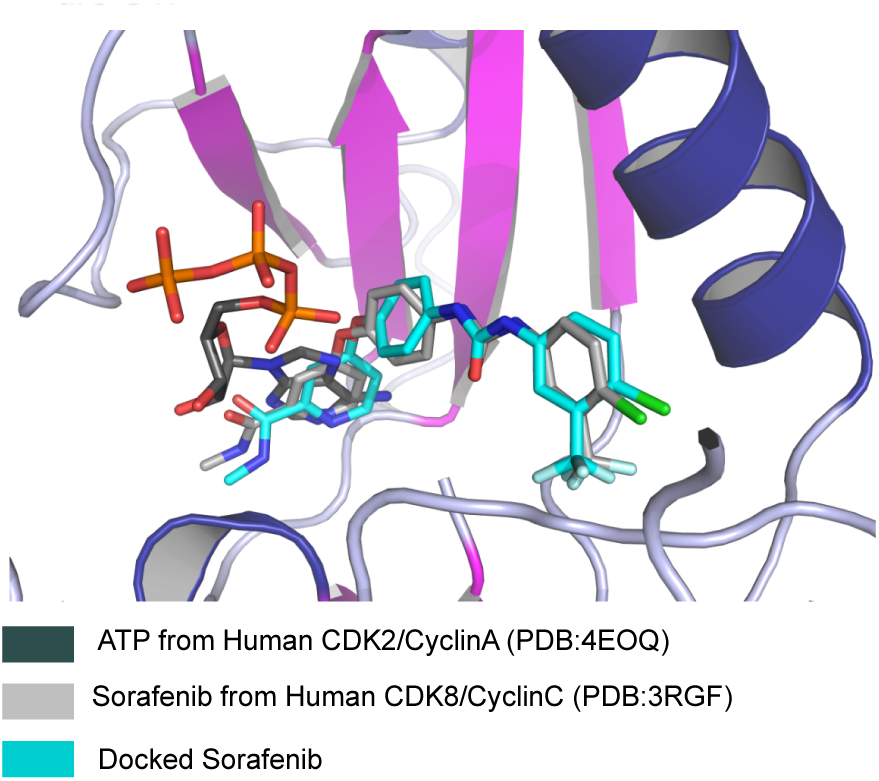
In silico screening model. Superposition of docked Sorafenib (cyan carbon atoms) onto its corresponding crystallographic structure. Sorafenib from 3RGF is depicted with light gray carbon atoms. As no structure of CDK8 with ATP is available, the ATP (dark gray carbon atoms) of the superposed structure of the human CDK2/Cyclin A complex (PDB ID: 4EOQ) is shown to highlight its potential binding site in CDK8.

**Fig. S2.**
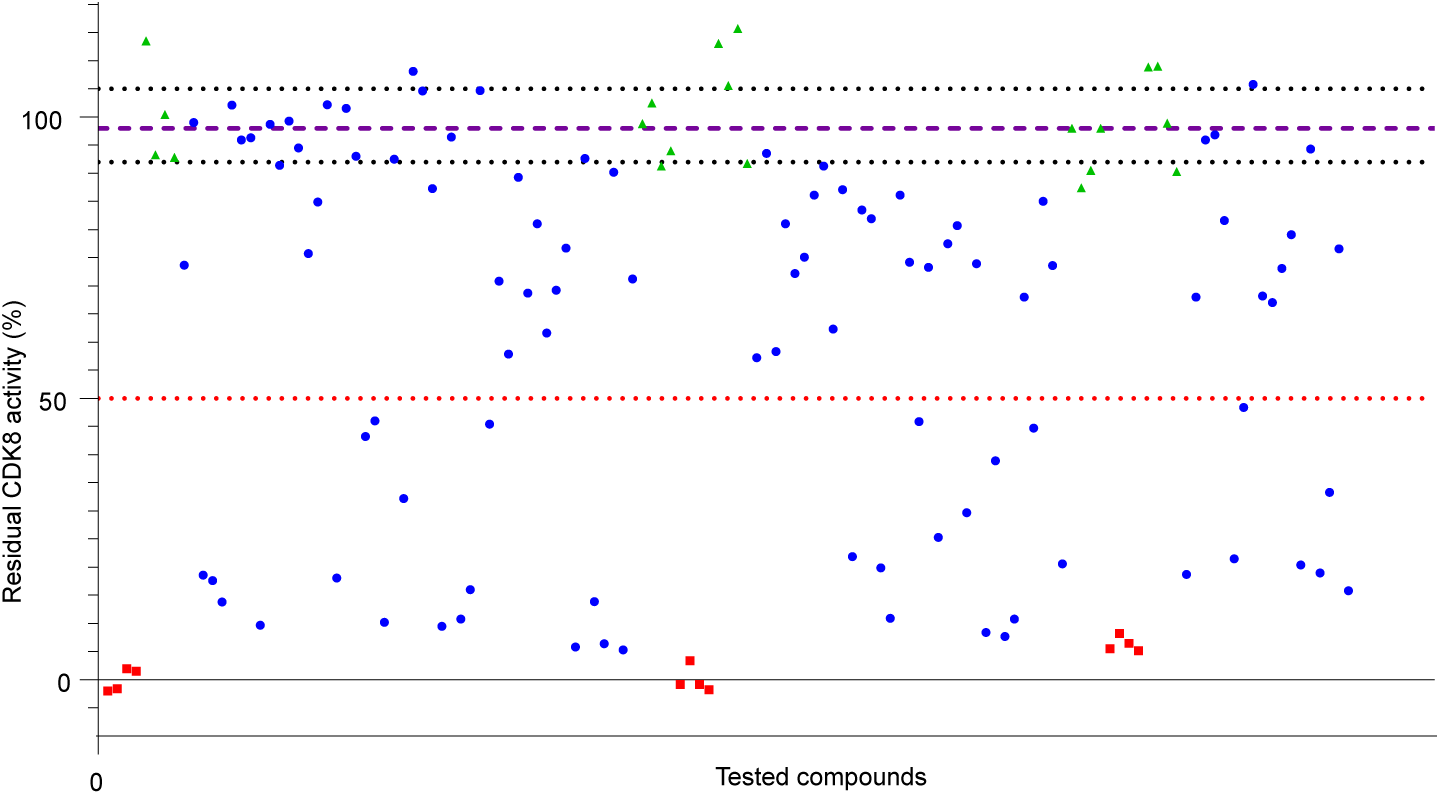
Primary assessment of in silico hits on CDK8 activity. Compounds were tested in duplicate at 10 µM in kinase assays with recombinant CDK8/Cyc C complex, in the presence of 15 µM [γ-^33^P] ATP. The graph represents the residual CDK8 activity (%) for each compound tested (blue dots); background controls (red dots) and negative control (DMSO 1%, green dots). 36 compounds inhibit CDK8 to below 50%.

**Fig. S3.**
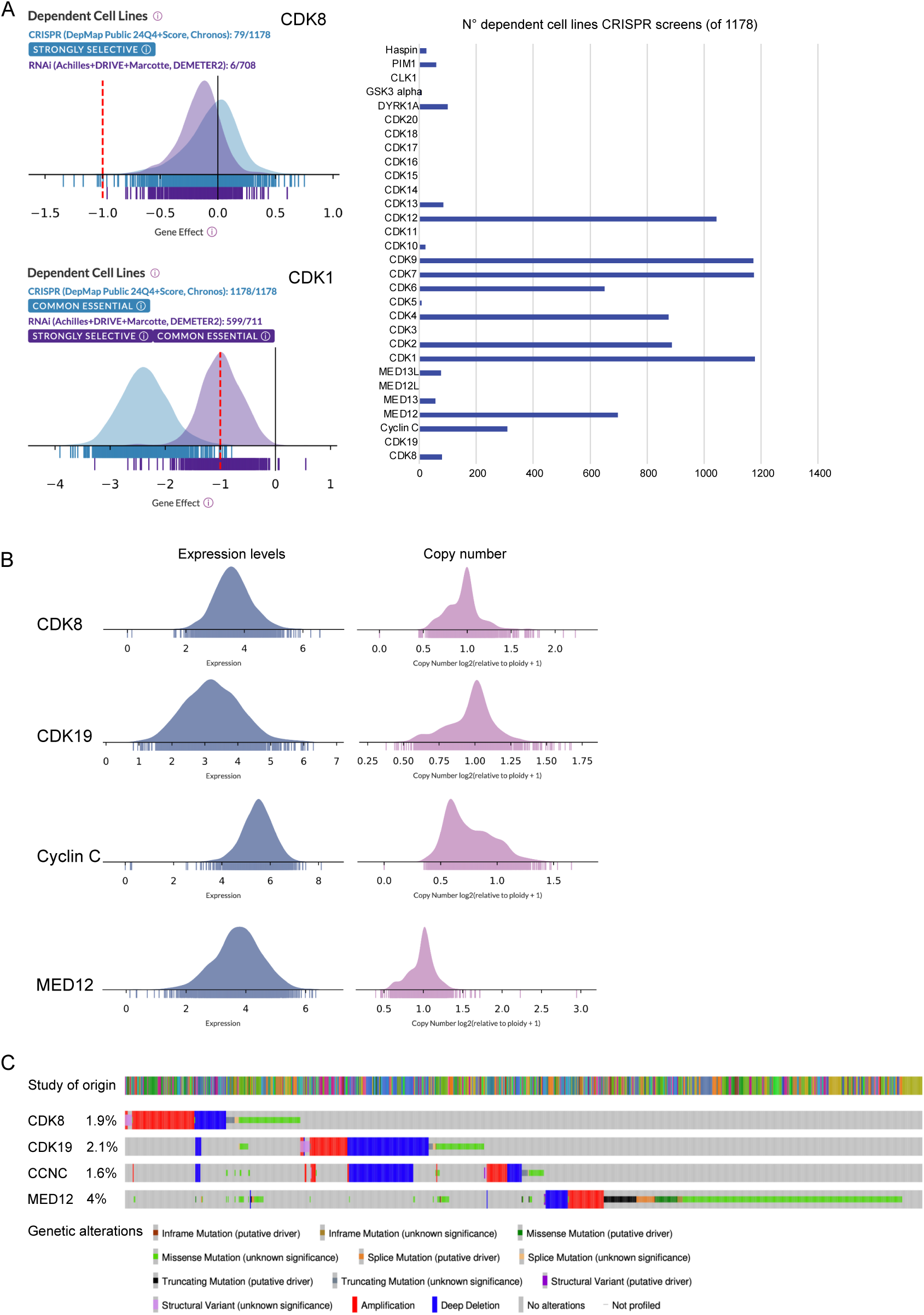
DepMap comparison of all CDK genes shows that CDK8 and CDK19 are non essential, while they are rarely mutated in human cancers. A. Left, DepMap analysis of dependecy of cancer cell lines on CDK8 and CDK1 (an example of an essential CDK), from genome-wide CRISPR screens. Score 0 indicates a non essential gene; score of -1 corresponds to a median of all common essential genes. Right, number of cancer cell lines dependent on the indicated gene according to the DepMap CRISPR screen database. All CDKs and kinases used to screen for CDK8/19 inhibitor specificity are shown. B. Expression levels (from RNAseq data, TPM-normalised and represented as log2(TPM+1)) and relative copy number (based on whole exome sequencing) in cancer cell lines. C. cBioPortal mutation analysis of the indicated genes in pancancer TCGA data.

**Fig. S4.**
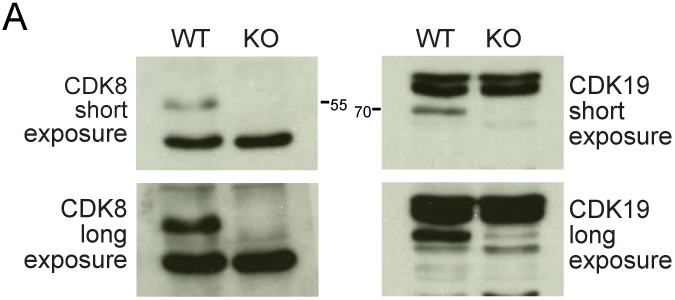
Knockout of CDK8 and CDK19 in SV40-immortalised MEFs. Verification of efficiency of knockout by Western blot (short and long exposures are shown).

